# Robust semi-supervised scRNA-seq integration from virtual adversarial learning

**DOI:** 10.64898/2026.06.07.730754

**Authors:** Chuan He, Paraskevas Filippidis, Jian Xing, Steven Kleinstein, Leying Guan

## Abstract

Single-cell RNA sequencing integration methods that rely solely on transcriptomic data often struggle to preserve fine-grained distinctions between closely related cell subtypes. As a result, cell populations that are separable in the raw data may become over-mixed after integration, reducing biological resolution and interpretability. Incorporating marker gene information can potentially address these issues; however, the variability and complexity of available marker sets limit their effective application. To address this, we introduce scCRAFT+, a semi-supervised integration model that innovatively incorporates marker gene information through Virtual Adversarial Training (VAT). By jointly optimizing marker-derived supervision and transcriptome-wide representations, VAT enforces local prediction smoothness among transcriptionally similar cells, improving robustness to noisy marker annotations while enhancing both integration quality and cell type auto-annotation. This targeted approach significantly enhances annotation accuracy and robustness, particularly when faced with incomplete or incorrect marker gene sets. Benchmarking shows that scCRAFT+ achieves consistently stronger performance than current unsupervised and supervised integration approaches, resulting in improved integration quality and biologically meaningful sub-cell type auto-annotations.

## 1 Introduction

The growth of single-cell RNA sequencing (scRNA-seq) has generated expansive datasets that reveal cellular diversity at an unprecedented scale [1–3]. However, most studies still sample relatively few individuals and exhibit study-specific biases due to differences in cohorts, sample handling, and technologies. Although directly pooling datasets could in principle increase statistical power, such naïve aggregation often amplifies technical artifacts and obscures true biological variation. To mitigate confounding batch effects in scRNA-seq integration, a landscape of unsupervised computational methods has emerged. Foundational tools, including Seurat[4], scVI[5], and Harmony[6], have seen wide adoption. Subsequent methods, such as Scanoroma[7] and scCRAFT[8], were developed to further enhance integration performance. While these unsupervised approaches effectively resolve transcriptional patterns of major cell lineages, achieving fine-grained resolution remains a challenge[4]. This limitation is particularly acute for immune lineages; for example, T cell subsets such as naïve versus central memory states, effector states, and *γδ* T cells are difficult to differentiate. based solely on unsupervised clustering of transcriptomic data without additional guidance [9, 10].

Resolving fine-grained cell subtypes often requires incorporating prior biological knowledge to guide the integration [11]. Well-curated marker gene sets represent a cost-effective and accessible source of this prior knowledge [11–15]. Many marker sets have been derived from large-scale, well-curated atlas data, providing comprehensive insights into cell-type definitions and disease-specific states from multiple sources [11, 15–18]. Others, such as those in Azimuth, leverage refined clustering from multi-omics approaches that incorporate surface protein expression to capture a more complete view of cellular states [4, 19–21]. Because of their high biological specificity, these markers offer an accessible solution for sub-cell type auto-annotation [12, 22–25], which can also serve as auxiliary information to improve data harmonization quality through supervised/semi-supervised integration [13, 26–29]. Public databases, including PanglaoDB [30] and CellMarker [31], further promote consistency and reproducibility.

Nevertheless, marker-based approaches face several significant challenges that underline their reliability. First, marker reliability is highly context-dependent, with performance varying substantially across datasets and databases [25, 32]. Second, many marker lists were designed for manual, hierarchical annotation workflows that allow flexible adjustment of cell-type nomenclature and resolution [11, 17, 18, 33]. Third, unsatisfying specificity of marker genes for certain cell types combined with high dropout rates in scRNA-seq data increases vulnerability to errors by relying solely on limited marker expression without accounting for broader patterns [25]. Our recent benchmark study further showed that leading semisupervised methods offered minimal gains over unsupervised approaches under realistic label-imperfection settings, including when using auto-annotated labels from state-of-the-art annotation tools [34]. Thus, although marker-derived labels can potentially provide useful biological guidance, we still lack an effective mechanism for extracting reliable information from such labels while remaining robust to their imperfections.

Here, we design a tailored Virtual Adversarial Training (VAT) module to robustly and efficiently integrate marker information into the learning process [35]. We combine this module with scCRAFT [8], which was previously shown to outperform other leading integration tools in extensive numerical experiments, to build scCRAFT+, a semi-supervised scRNA-seq integration and label annotation tool. scCRAFT+ is designed to tolerate incorrect labels and imperfect marker lists while efficiently leveraging reliable label information. Using publicly available marker data, scCRAFT+ surpasses state-of-the-art unsupervised and semi-supervised methods and produces high-resolution cell-type annotations.

## 2 Results

### 2.1 Overview of scCRAFT+

A key challenge in high-resolution cell type annotation is that available labels are frequently incomplete, noisy, or overconfident, making it difficult to balance robustness and annotation efficiency without dataset-specific tuning. scCRAFT+ addresses this by extending the scCRAFT framework [8] with a marker-guided virtual adversarial training (VAT) module that simultaneously exploits fine-grained marker information and confers resilience to label noise (Fig. 1).

**Fig. 1.**
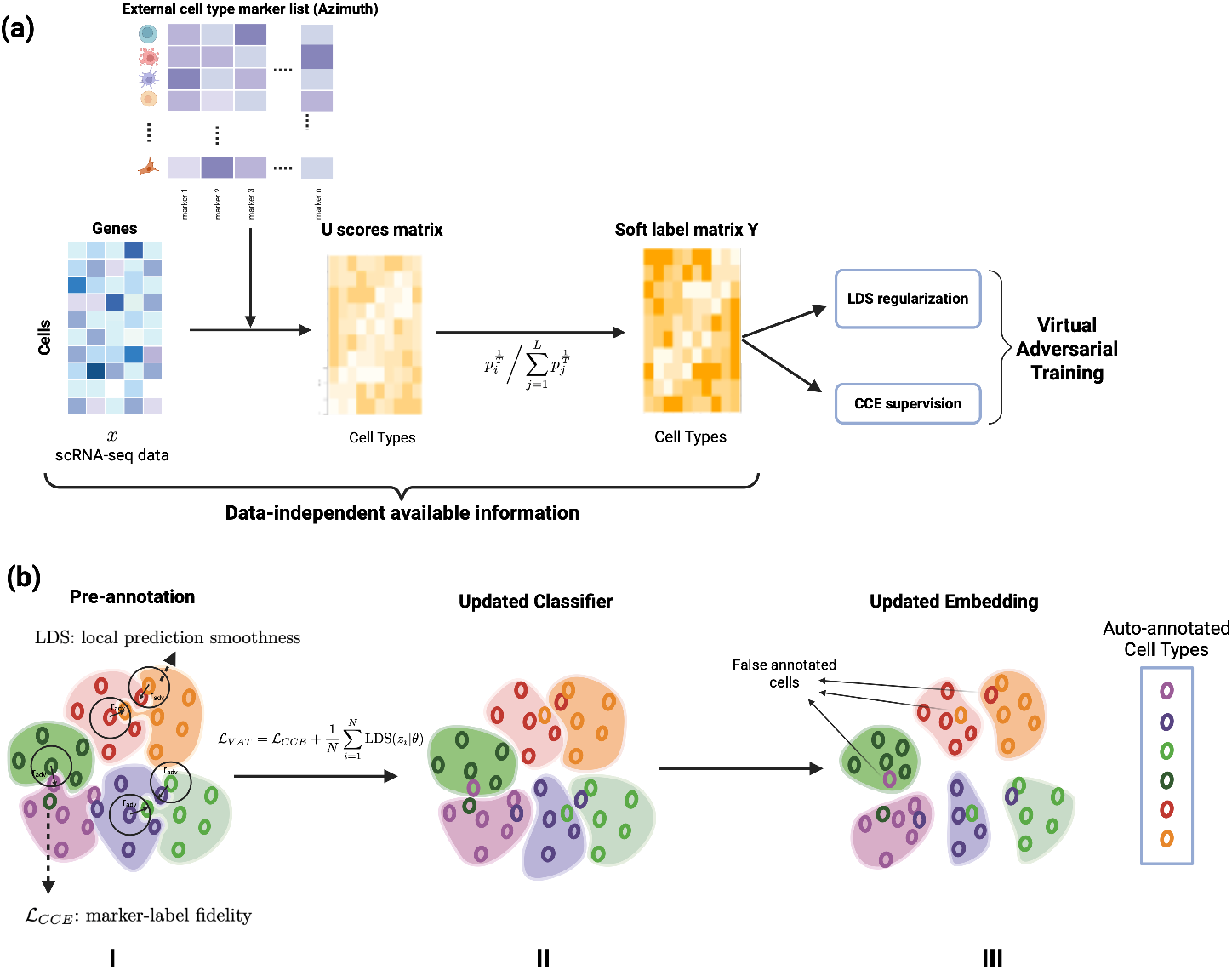
Overview of scCRAFT+ (a) Schematic diagram of the marker-guided VAT module. Given an external marker list, scCRAFT+ computes U scores from ranked marker expression and sharpens them into a marker-derived soft label matrix *Y*. The soft labels provide weak supervision through CCE, while LDS regularization promotes locally smooth predictions in the latent space. (b) Illustration of the VAT-guided learning process. Panel I shows marker-derived pre-labeling and the combined VAT objective. Panel II shows VAT-smoothed classifier predictions within transcriptionally coherent neighborhoods. Panel III shows the resulting VAT-guided embedding and auto-annotation, where scattered false labels are reduced while major cell type structure is preserved.

scCRAFT+ processes gene expression data *X* through the variational autoencoder (VAE) backbone of scCRAFT to generate a low-dimensional embedding *Z* = *E*_*θ*_(*X*) that captures the underlying transcriptional structure. This embedding is passed to an annotation classifier guided by a marker-derived cell type assignment vector *Y ∈* [0, 1]^*K*^, constructed via a fast U-score computation that summarizes ranked marker expression across *K* cell types (Fig. 1a). Rather than relying on a standard cross-entropy objective, classifier optimization employs a VAT training objective [35] comprising a cross-entropy (CCE) loss over the labeled cell set 𝒱and a local distributional smoothness (LDS) regularization term computed over all *N* cells (Fig. 1b):

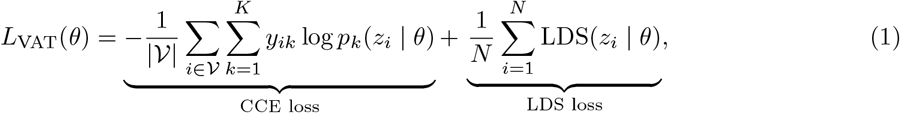

where *y*_*ik*_ is the auto-annotated label assignment for cell *i* and type *k, p*_*k*_(*z*_*i*_ | *θ*) is the predicted probability from scCRAFT+, and LDS(*z*_*i*_ | *θ*) penalizes large prediction shifts under a virtual adversarial perturbation *r*_*adv*_ of the latent representation *z*_*i*_. The full model is then trained by jointly minimizing *L*_scCRAFT+_ = *L*_scCRAFT_(*θ*) + *λ*_*C*_ *L*_VAT_(*θ*), with *λ*_*C*_ = 1 by default. Definitions of *L*_scCRAFT_, the U score construction and full implementation details are provided in the Methods (Section 4).

Unlike standard cross-entropy training, which is sensitive to misspecified or overconfident labels, the LDS term in Eq. (1) provides implicit regularization against label noise by encouraging locally consistent predictions in the latent space. This design allows scCRAFT+ to exploit high-resolution, marker-derived annotations for fine-grained cell type discrimination while remaining robust to annotation errors that are common at high resolution (see Methods).

### 2.2 scCRAFT+ is robust to incomplete and noisy annotations

We first benchmarked scCRAFT+ on four real-world datasets (PBMC, Pancreas, Lung atlas, and Human Immune) (Supplementary Section 1), under controlled incomplete and erroneous cell type label settings. We generated progressively challenging scenarios by introducing 0%–70% of either missing or incorrect cell type labels. We compared scCRAFT+ against three leading semi-supervised methods (scANVI [26], scGen [29], and ssSTACAS [13]) alongside their respective unsupervised counterparts (scVI for scANVI and scGen; scCRAFT for scCRAFT+; Seurat RPCA for ssSTACAS) (Supplementary Section 2). All simulated labels, including incorrect ones, were provided as labels to assess robustness under realistic conditions. Integration accuracy was evaluated using two composite scores: a bioconservation score averaging NMI, ARI, and ASW cell, and a batch-mixing score averaging ASW batch, Graph Connectivity, kBET, bLISI, and True Positive Rate (Supplementary Section 3).

In our first setting, we assessed robustness and efficacy in utilizing incomplete labels by varying missing annotations from 0% (fully annotated) to 70%. scCRAFT+ demonstrated superior performance, particularly in batch mixing, where it was the only semi-supervised method to consistently improve over its unsupervised counterpart at all levels of label availability (Fig. 2a). Among the competing methods, scGen matched scCRAFT+ in bio-conservation under complete labels but deteriorated sharply as missing annotations increased. ssSTACAS was stable across missingness levels yet provided negligible gains over its Seurat baseline on most datasets, suggesting limited benefit from its semi-supervision component. scANVI improved over scVI in bio-conservation but remained consistently inferior to scCRAFT+ across all datasets, with a particularly large gap on the Human Immune dataset; its batch-mixing performance lagged behind both scCRAFT+ and scVI.

**Fig. 2.**
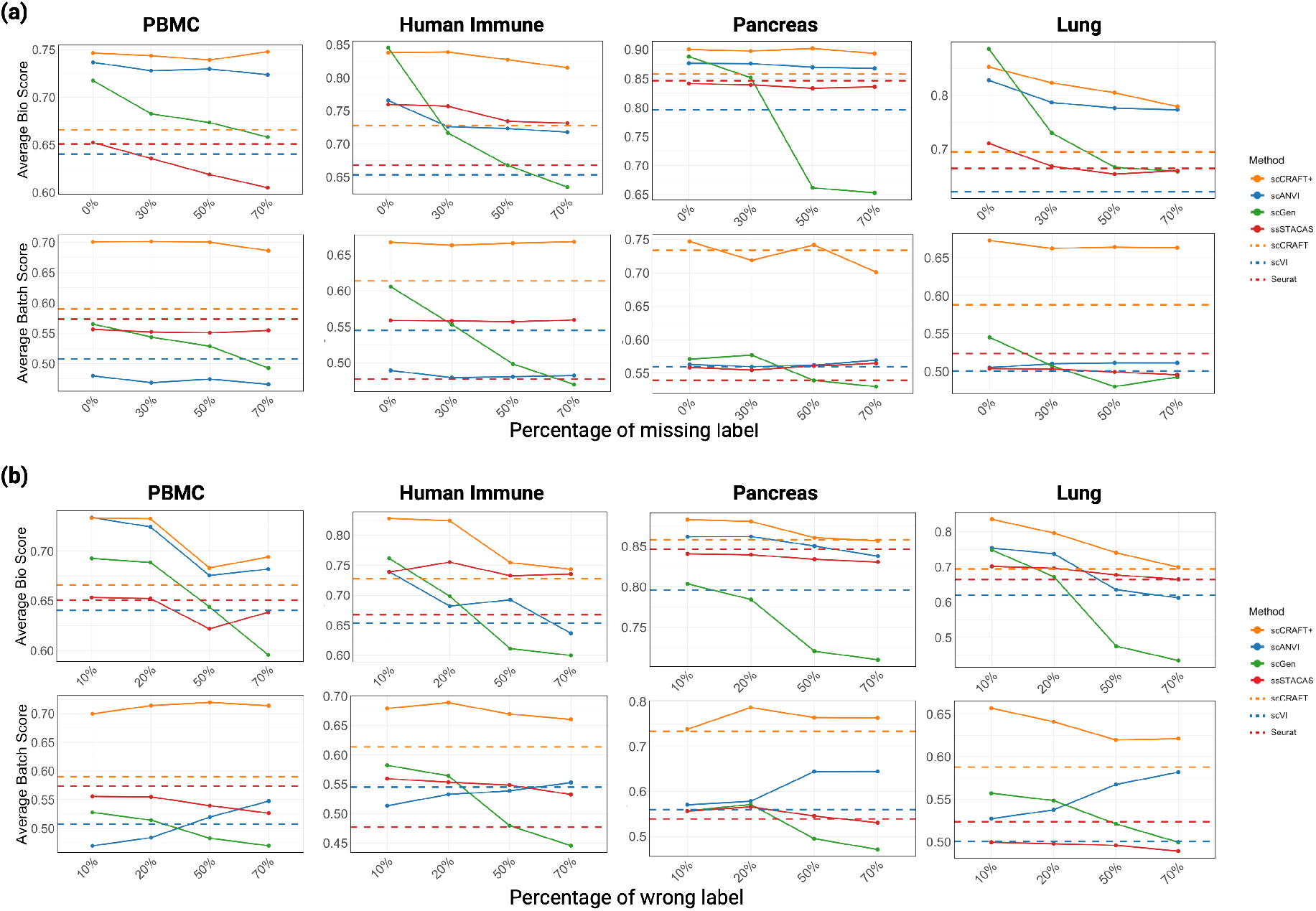
Benchmarking on integration performance of supervised and semi-supervised methods under incomplete and noisy cell type annotations settings (a) Preservation of biological variance (measured by averaging three bio-conservation metrics) and batch mixing (measured by averaging five batch-mixing metrics) for four data integration tasks, using full cell type labels (0% missing) as input and increasing the percentage of missing cell type labels (up to 70%). (b) Preservation of biological variance (measured by average biological metrics) and batch mixing (measured by average batch correction metrics) for 4 data integration tasks, using as partially wrong labels in all known labels (10%) and increasing the percentage of wrong cell type labels (10% to 70%). Unsupervised versions of scCRAFT+, ssSTACAS and scANVI (scCRAFT, seurat and scVI respectively) are included for reference. Source data is provided as a Source Data file.

In the second setting, we assessed robustness to label noise by varying the mislabeling proportion from 10% to 70%. scCRAFT+ consistently outperformed its unsupervised counterpart scCRAFT across all datasets in both bio-conservation and batch-mixing even at 70% label corruption, and exhibited a substantial improvement in batch-mixing over all semi-supervised baselines (Fig. 2b), underscoring its capacity to extract useful signal from substantially inaccurate annotations. In terms of bio-conservation, ssSTACAS exhibited the most stable performance as mislabeling increased, followed by scCRAFT+ and scANVI, with scGen showing the greatest deterioration. Despite this stability, ssSTACAS consistently achieved lower bio-conservation scores than scCRAFT+ across all mislabeling levels, with minimal gain over its unsupervised counterpart Seurat except on the Human Immune dataset — consistent with the pattern observed in the first setting. Regarding batch-mixing, scANVI showed a distinct and consistent increase across all datasets as the mislabeling proportion grew, indicating that label noise induced excessive inter-batch mixing without a corresponding improvement in biological structure preservation

Together, these results highlight scCRAFT+’s high label utilization efficiency, achieving the best overall integration performance by robustly leveraging partially available or correct annotations.

### 2.3 Automated marker-based pre-annotation enhances integration performance of scCRAFT+

We next assessed integration quality across semi-supervised methods when cell type labels are derived from marker-based auto-annotation. To ensure concistency, we generated auto-annotations for all methods using fine-grained marker lists from the corresponding tissue in the well-curated Azimuth atlas [4], combined with the fast U score based auto-annotations (see Methods). Azimuth was selected as the annotation reference because it was observed to yield the highest-quality auto-annotated labels for guiding semi-supervised integration in a systematic comparison against marker-based pipelines SingleR and CellAssign, as well as the deep learning strategy TOSICA, in out-of-sample generalization benchmarks [34].

scCRAFT+ outperformed alternative methods across four integration tasks in both bio-conservation and batch mixing (Fig. 3a, Supplementary Fig. S6-S9 in Supplementary Section 7), with its unsupervised predecessor scCRAFT ranking second overall. scCRAFT+ improved consistently over scCRAFT across all datasets, though the magnitude of the gain varied: the PBMC dataset showed the largest improvement, with a 5.7% increase in bio-conservation and a 7.0% enhancement in batch mixing. Although scANVI achieved a 5.6% collective improvement in bio-conservation over its unsupervised counterpart scVI, its batch-mixing score was collectively 2% lower than scVI, indicating a trade-off between biological and technical variation removal under auto-annotated labels. Consistent with our controlled experiments, ssSTACAS was stable but provided minimal gain over Seurat except on the Human Immune dataset, where a collective 1.5% improvement in bio-conservation was observed. scGen was frequently inferior to its unsupervised counterpart scVI under auto-annotated labels, consistent with its high sensitivity to label quality observed in the controlled noise setting. Notably, scANVI, ssSTACAS and scGEN all lagged behind scCRAFT+ by a substantial margin.

**Fig. 3.**
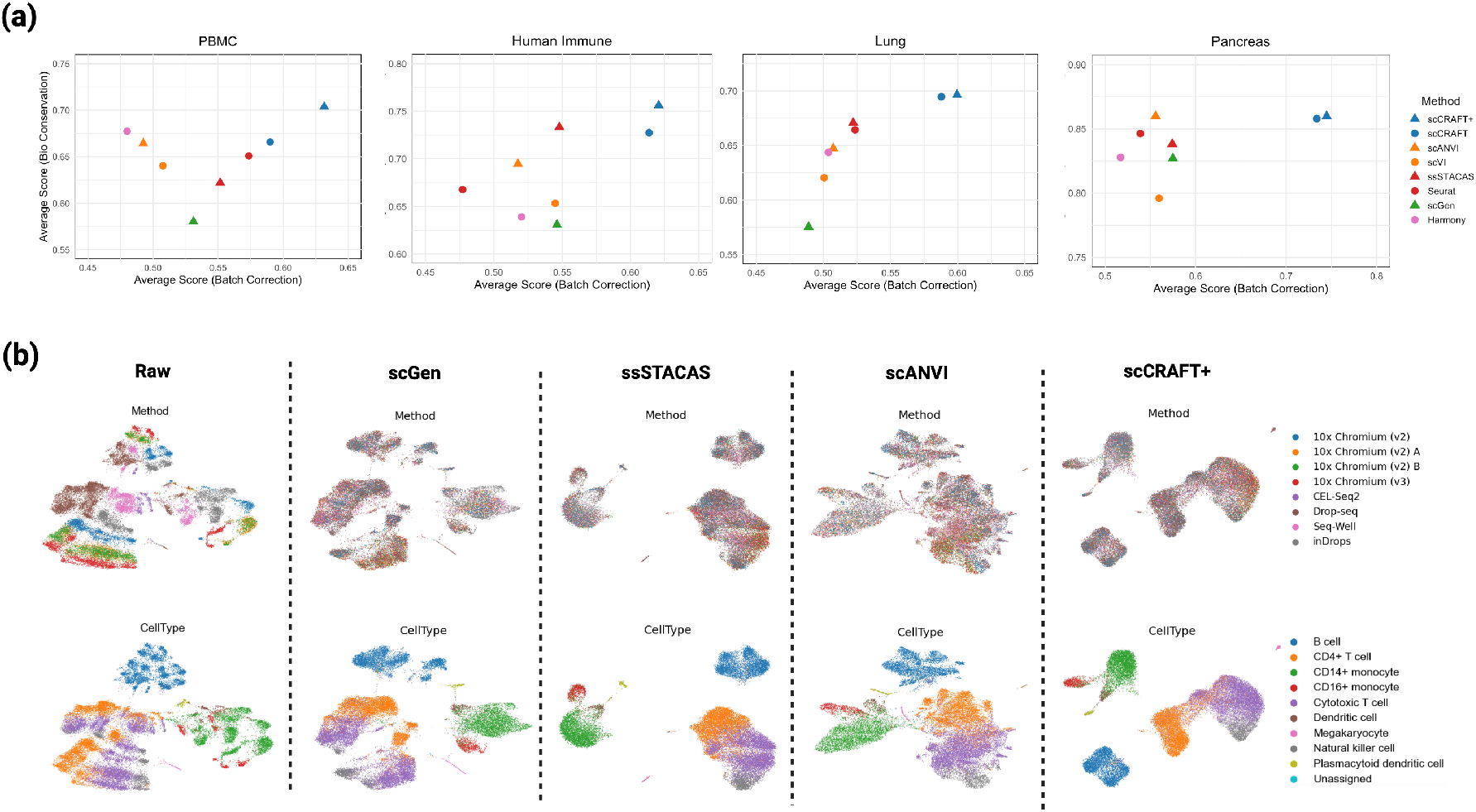
Integration performance for integration methods over 4 different tasks given auto-annotated cell type labels. (a) Scatterplot of average bio-conservation score against the average batch mixing score for each method. (b) UMAP visualization of the PBMC dataset, depicting the unintegrated raw data and embeddings from four semi-supervised integrated methods colored by batches and cell types.

Focusing on the PBMC dataset, where scCRAFT+ showed the largest gain over scCRAFT, a visual inspection of the embedding confirmed that only scCRAFT+, scCRAFT, and Harmony achieved clear clustering and separation of lymphoid immune cell populations without visible fragmentation of the same cell type across batches (Fig. 3b, Supplementary Fig. S6). Among these, scCRAFT+ better resolved CD14+ monocytes, CD16+ monocytes and dendritic cells compared to scCRAFT and Harmony, with

Harmony’s batch-mixing score 31.5% lower than scCRAFT+, consistent with its improved quantitative metrics (Fig. 3a).

To isolate the contribution of the VAT mechanism from that of the integration backbone, we conducted ablation studies comparing supervision with and without the VAT mechanism. The results confirm that VAT enhances robustness to label overconfidence inherent, and that replacing the standard cross-entropy loss with the VAT objective improves integration performance for both scCRAFT+ and scANVI using auto-annotated labels (Supplementary Section 6).

### 2.4 scCRAFT+ is robust to wrong marker list

Cell type composition can vary substantially across tissues, and a given cell type may express tissuespecific gene markers due to causes such as tissue homing, migration, or residency. This variability poses a significant challenge in annotating cells from an understudied tissue where a lack of high-quality reference markers can be common (e.g., skin in the Azimuth dataset). Consequently, the low quality of available markers can further degrade the performance of semi-supervised methods, sometimes causing them to perform no better or even worse than their unsupervised counterparts. To evaluate how scCRAFT+ and other semi-supervised approaches address this critical challenge, we designed an experiment to test the effects of using a tissue-mismatched marker list on integration quality. We utilized cell type reference marker lists derived from pancreas, lung and PBMC as cell type reference markers for auto-annotation when evaluating the four datasets under consideration. For clarity, we refer to the tissue-aligned marker-dataset combination as the “right” scenario: using PBMC markers for annotating the PBMC and Human Immune datasets, lung markers for the Lung dataset, and Pancreas markers for the Pancreas dataset. Conversely, we refer to the results from using tissue-misaligned markers (the tissue-mismatched scenarios) by the tissue origin of the markers used.

Sensitivity to marker list accuracy varied across methods. Consistent with our earlier synthetic data experiments, ssSTACAS demonstrated the most stable performance using different marker lists (Fig. 4). However, its improvement over its unsupervised counterpart, Seurat, was limited, except for the Human Immune dataset, and was generally much worse than scCRAFT+ when using either the tissue-aligned “right”) or misaligned marker lists. scGen was highly dependent on accurate cell annotations, a finding also reflected in our synthetic data experiments. Recall that even with tissue-aligned auto-annotations, its performance was overall sub-optimal and did not outperform its unsupervised counterpart scVI. The misaligned tissues autoannotation further deteriorated its performance measured by the combined bio-conservation and batch-mixing score with a large performance reduction of 29.8% in the Pancreas dataset and 11.8% in the Lung dataset. Both scANVI and scCRAFT+ achieved relatively stable performance when varying the reference marker lists, placing their stability level between that of ssSTACAS and scGen. Unlike scCRAFT+, scANVI did not show improvement over its unsupervised counterpart scVI even when using decent-quality labels, as was demonstrated in both our synthetic data experiments and the auto-annotations based on “right” marker lists. Consequently, we focus on comparing the bio-conservation scores achieved by scANVI and scCRAFT+ using different auto-annotations. Overall, scANVI observed a 5.2% average drop in bio-conservation when comparing the tissue-aligned marker result to the poorest misaligned marker result, whereas scCRAFT+ observed only a 2.2% drop in the same comparison. Notably, in the Lung dataset, scANVI observed a large drop of 9.1% in bioconservation, changing from 4.3% better than scVI to being 4.6% worse than scVI, whereas scCRAFT+ maintained at least comparable performance levels to scCRAFT across the different marker lists. Consistent with this robustness, the cross-tissue annotation examples in Supplementary Section 5 show that mismatched marker lists can still provide coarse information for shared cell populations, while a high proportion of “Unknown” predictions can help users recognize that the marker reference may be poorly matched to the target tissue (Supplementary Section 5).

**Fig. 4.**
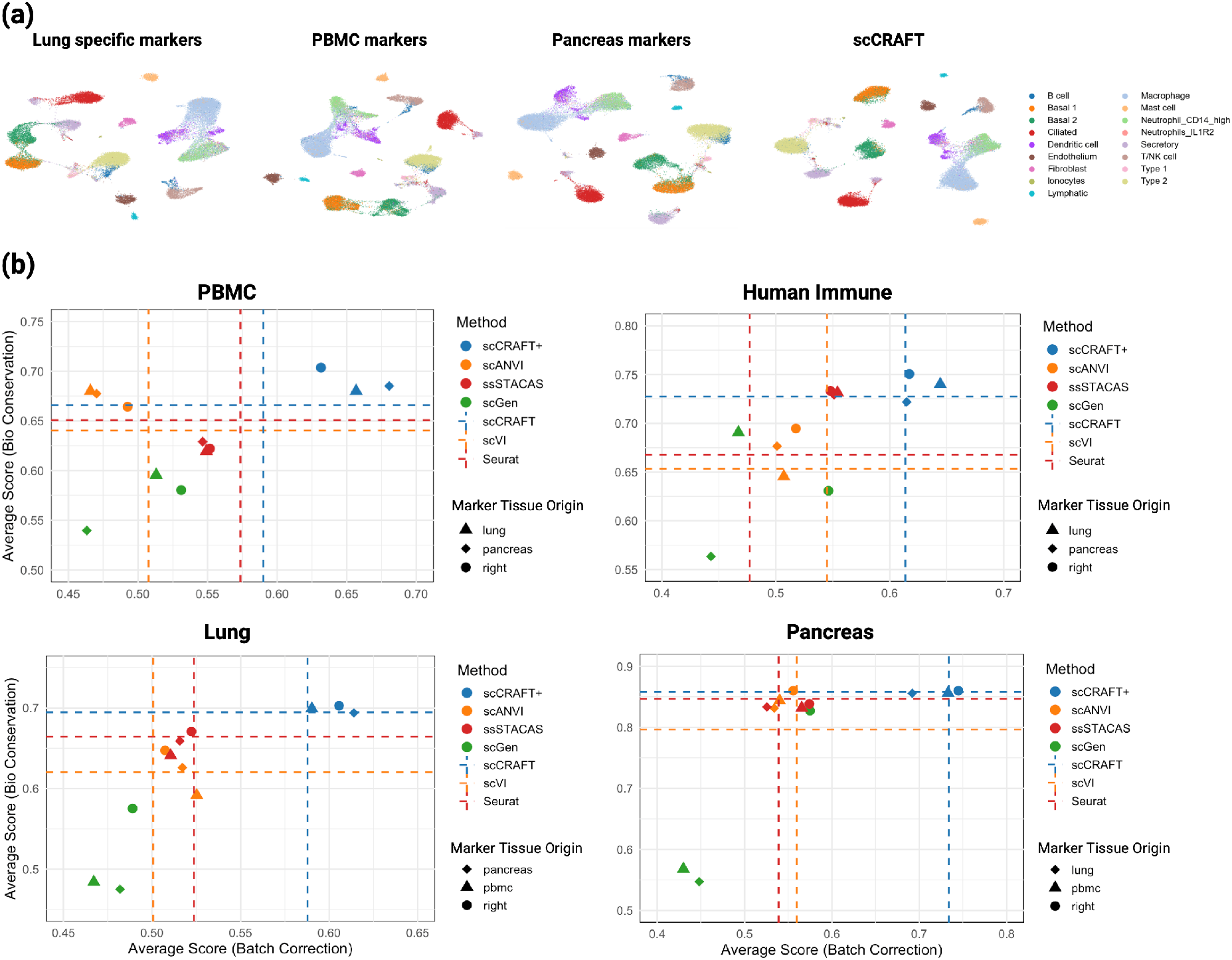
Performance comparison of semi-supervised integration methods using correct and mismatched marker lists. (a) UMAP visualization of lung dataset integration, depicting the scCRAFT+ integration performance when applying different sets of markers. (b) Scatterplot of the average bio-conservation score against the average batch mixing score, showing the performance of four semi-supervised methods across four integration tasks. The legend annotations indicate the marker set used for each task: “Right” refers to the correct dataset-specific markers, including the PBMC marker list level 3 for PBMC and human immune datasets, the HLCA level 2 markers for the lung atlas, and pancreas-specific markers for the pancreas dataset. “Lung” refers to the use of lung-specific markers applied to unrelated datasets, while “Pancreas” and “PBMC” represent the use of pancreas and PBMC markers, respectively, in non-matching datasets.

### 2.5 scCRAFT+ provides reliable and high-resolution cell type annotation

High-resolution cell type annotation requires distinguishing closely related subtypes while avoiding unsupported fine-grained assignments. While comprehensive reference atlases facilitate the annotation of rare cellular subsets, high-resolution mapping introduces the risk of over-annotation and spurious assignment. In practice, this may split a transcriptionally coherent cluster into multiple weakly supported labels or introduce mosaic labels from unrelated lineages. scCRAFT+ can successfully mitigate this dilemma through VAT-mediated manifold regularization. To support this, we compared the source-provided annotations, scCRAFT+ annotations, and Azimuth predictions [4] for the Human Immune Dataset. For a controlled comparison, both scCRAFT+ and Azimuth used the same Azimuth Level III PBMC marker reference. Relative to the source annotations, scCRAFT+ clearly separated major myeloid, lymphoid, and non-immune populations (Fig. 5a) and, in some cases, yielded more specific labels as a result of the broader coverage of the reference. Examination of marker genes associated with these predicted cell types further supported the validity of scCRAFT+ predictions (Fig. 5b).

**Fig. 5.**
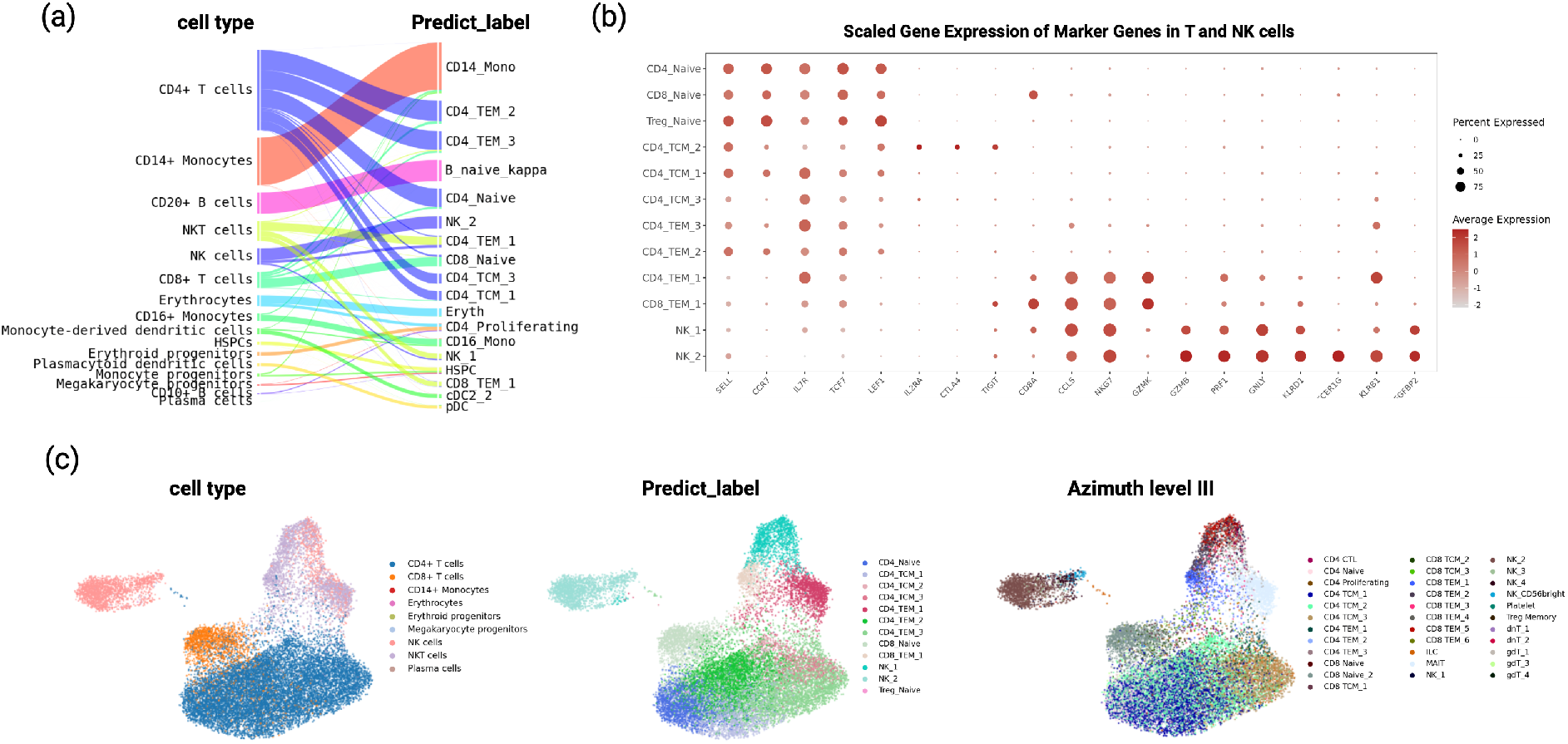
Benchmarking scCRAFT+ for automated cell type annotation on human immune data. (a) Alluvial plot of the entire human immune dataset comparing the original label and the predicted label given by scCRAFT+ (b) Average gene expression of T and NK cell marker genes in the predicted cell subtypes. (c) UMAP visualization of scCRAFT+ embedding of T and NK cells in the Human Immune dataset, with the color labels as cell types (original), predicted label (given by scCRAFT+) and Azimuth level III auto-annotation.

We next compared scCRAFT+ and Azimuth on the finer-grained task of T cell subtype annotation, where high diversity makes reference-based labeling particularly challenging. Within the T cell compartment, Azimuth assigned 34 of 38 reference labels, including 12 CD8 T cell subtypes, but these labels showed substantial overlap and weak separation. In contrast, scCRAFT+ assigned 11 of 38 labels and produced more distinct subtype boundaries, avoiding labels not supported by the data (Fig. 5b, c).

## 3 Discussion

Here, we introduce scCRAFT+, a semi-supervised method for integrating single-cell RNA-seq data with help from high-resolution marker-based cell type information. Marker labels are useful, but they are often incomplete, noisy, or only partially suited to the dataset being analyzed, especially when we want to utilize high-resolution information from such prior knowledge beyond broad cell lineages. scCRAFT+ uses Virtual Adversarial Training to make efficient use of these labels while remaining robust to the aforementioned imperfection.

scCRAFT+ showed consistently strong integration performance across multiple label-guided settings. Under given-label supervision, it remained robust to both missing and incorrect annotations, outperforming existing semi-supervised methods in biological conservation and batch correction. When using marker-derived auto-annotations, scCRAFT+ further improved over its unsupervised counterpart, indicating that the framework can effectively convert imperfect marker information into useful integration guidance. Its stable performance under mismatched marker lists further suggests that scCRAFT+ can extract transferable biological signals from partially overlapping references while avoiding substantial degradation when marker information is incomplete or tissue-misaligned. Beyond integration, scCRAFT+ provides reliable high-resolution cell type annotation. In the Human Immune dataset, scCRAFT+ generated annotations that were more coherent with the transcriptomic embedding structure than direct Azimuth annotation, particularly within T and NK cell compartments. Marker expression analysis further supported the predicted lymphoid annotations, demonstrating that scCRAFT+ can use marker information to guide fine-grained annotation without forcing all cells into overly detailed reference categories.

Still, the final naming of closely related cell states should require manual inspection. scCRAFT+ combines marker-derived information with cell representations learned from gene expression, which helps keep labels tied to real transcriptional separation. However, for closely related populations, there may still be no single obvious name. The input marker database may also reflect one annotation convention rather than a community-wide agreement. For example, in the Human Immune dataset, one cluster predicted as NK1 overlapped with source annotations containing both NK and NKT cells, while Azimuth labeled similar cells as CD8 TEM. This type of disagreement is not only a model limitation. It also reflects the fact that computational methods can learn structure, but cell type names still depend on the reference definitions provided by researchers.

One limitation of this study is that the quantitative benchmarks rely on source-provided annotations as the ground truth, but these annotations are themselves often derived from unsupervised clustering followed by manual marker inspection, which may contain errors themselves. In addition, source annotations are often less detailed than the marker references used for auto-annotation. This can make it harder for the metrics to capture whether a method has placed refined subtypes correctly. Despite this limitation, author-provided labels remain one of the most practical and transparent choices for benchmarking.

Overall, scCRAFT+ provides a practical way for robust semi-supervised scRNA-seq integration and marker-guided annotation. Its main contribution is to use VAT to transform imperfect marker-derived supervision into stable, biologically coherent embeddings and annotations.

## 4 Methods

### 4.1 Data preprocessing

All scRNA-seq datasets were processed with the same Scanpy workflow [36]. Raw count matrices were loaded as AnnData objects. Cells expressing fewer than 300 genes and genes detected in fewer than five cells were removed. Library sizes were normalized to 10,000 counts per cell and then transformed with log(1 + *x*). For each integration task, we selected the top 2,000 highly variable genes while accounting for batch structure. The same preprocessing steps were applied before running scCRAFT+ and all benchmarked methods.

### 4.2 scCRAFT+ model

Let *X ∈* ℝ^*N×G*^ denote the processed expression matrix for *N* cells and *G* genes, and let *b*_*i*_ denote the batch label for cell *i*. scCRAFT+ builds on the scCRAFT backbone [8]. The backbone contains an encoder *E*_*θ*_, a decoder *G*_*ϕ*_, and a batch discriminator *D*_*ψ*_. The encoder maps each expression vector *x*_*i*_ to a latent representation *z*_*i*_ = *E*_*θ*_(*x*_*i*_), giving the embedding matrix *Z* = *E*_*θ*_(*X*). The decoder reconstructs the input expression profile from *z*_*i*_, while the discriminator predicts the batch label from *z*_*i*_. Here, we give a brief recap of the used scCRAFT backbone for the sake of completeness. The VAE component is trained with a modified negative evidence lower bound,

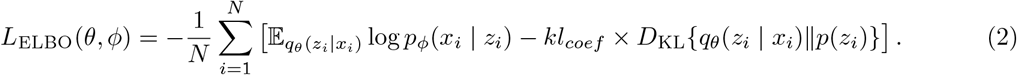

The reconstruction is further regularized by a cosine loss between the observed and reconstructed expression profiles,

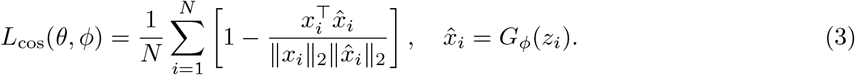

Batch correction is implemented through adversarial domain adaptation. The discriminator is trained to predict the batch label from the latent embedding,

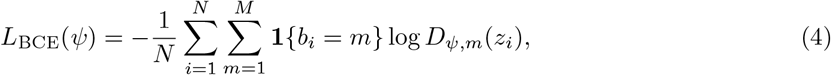

where *M* is the number of batches and *D*_*ψ,m*_(*z*_*i*_) is the discriminator probability for batch *m*. During encoder training, this term is reversed so that *E*_*θ*_ learns embeddings that are less predictive of batch.

To preserve biological structure, scCRAFT also uses a dual-resolution triplet loss for biological topology preservation within the same batch. For a triplet (*a, p, n*) in a given batch, the anchor cell *a* and positive cell *p* are selected from the same high-resolution biological neighborhood or clustering group, while the negative cell *n* is selected from a different low-resolution clustering group. The loss is

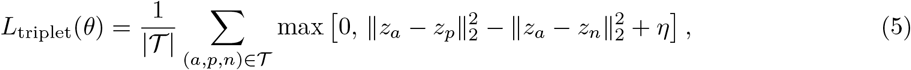

where *η* is the triplet margin and 𝒯 contains triplets sampled across two clustering resolutions. This term encourages cells with similar transcriptomic structure to remain close while separating cells from different neighborhoods.

The scCRAFT backbone objective optimized by the encoder and decoder is

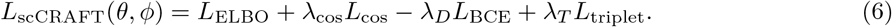

The negative sign before *L*_BCE_ reflects the adversarial update for the encoder. The discriminator itself is trained to minimize *L*_BCE_. We used the default scCRAFT weights in all benchmarking experiments: *kl*_*coef*_ = 0.005, *λ*_cos_ = 20, *λ*_*D*_ = 0.2 and *λ*_*T*_ = 1.

scCRAFT+ extends this backbone by adding a VAT-guided cell type classifier *C*_*ω*_ to the latent embedding. The classifier outputs *p*_*k*_(*z*_*i*_ | *ω*), the predicted probability that cell *i* belongs to cell type *k*, for *k* = 1, …, *K*. Label information enters the model as a matrix *Y ∈* [0, 1]^*N×K*^, where *y*_*ik*_ is the supplied weight for assigning cell *i* to type *k. Y* may be derived from source annotations, manual annotation, marker-based auto-annotation or another external annotation procedure. Thus, the Fast U score procedure described below is an input-label construction step used in this study, rather than part of the core scCRAFT+ model.

The full scCRAFT+ objective is

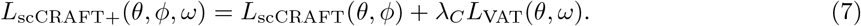

We used *λ*_*C*_ = 1 for the VAT-guided label objective.

### 4.3 VAT-guided use of label information

*V V*

The VAT-guided classifier is trained with a supervised label term and an unsupervised local smoothness term. Let 𝒱 be the set of cells with available label information in *Y*. For cells in 𝒱, the cell type cross-entropy (CCE) loss is

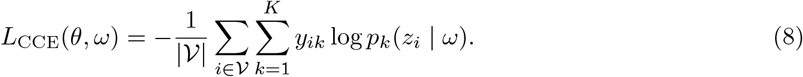

This term uses the supplied labels as guidance, but does not require them to be perfectly certain or perfectly correct.

The LDS term is applied to all cells and penalizes abrupt prediction changes near each latent embedding. For cell *i*,

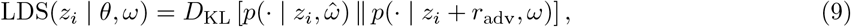

where *D*_KL_ is the Kullback–Leibler divergence and *ŵ* denotes a stop-gradient copy of the current classifier parameters. The perturbation *r*_adv_ is chosen as the direction within an *ϵ*-ball that most changes the classifier output:

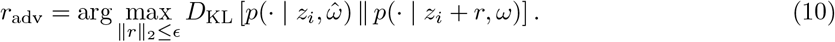

Following standard VAT [35], *r*_adv_ was approximated by power iteration. The LDS loss was averaged over all cells:

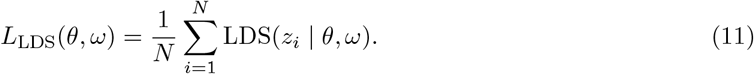

The VAT label objective is

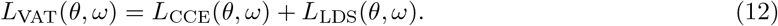

The perturbation radius was set to *ϵ* = 500, based on the sensitivity analysis in Supplementary Section 6.4.

### 4.4 Construction of marker-derived labels for benchmarking

For experiments using marker-guided auto-annotation, we constructed *Y* from external marker lists before fitting scCRAFT+. This step was used to provide a consistent label source across methods. It is separate from the scCRAFT+ core implementation, which only requires an input label matrix *Y*.

For each tissue, we used the corresponding Azimuth marker reference when available and selected the most detailed annotation level used in the benchmarking experiments. Marker lists can also be manually adjusted when a study has a more appropriate tissue-specific or disease-specific reference.

For each candidate cell type *k*, let *S*_*k*_ denote its marker set and *n*_*k*_ = |*S*_*k*_|. We used a fast rank-based score following the UCell strategy [24]. For each cell *i*, genes were ranked by log-normalized expression, and *R*_*ij*_ denotes the rank of gene *j* in that cell. The raw score for cell *i* and type *k* was

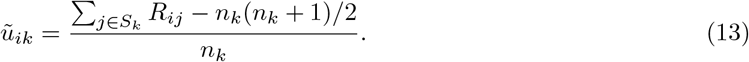

Raw scores were rescaled within each cell across the *K* candidate types:

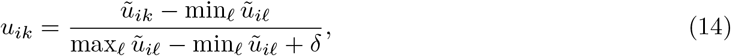

where *δ* is a small constant used only to avoid division by zero. The label matrix used in the autoannotation experiments was then constructed by temperature sharpening,

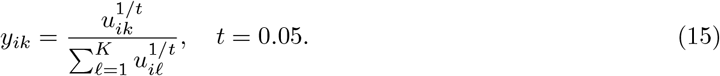

We used *t* = 0.05 as the default throughout the benchmarking experiments. This value gives confident marker-derived labels while retaining a probability vector over candidate cell types. Importantly, the method’s robustness to label overconfidence comes from the VAT objective rather than from fine-tuning this temperature. In sensitivity analyses, scCRAFT+ with VAT maintained similar integration quality under *t* = 0.05 and near-hard assignment (*t* = 0), whereas the cross-entropy-only variant became more sensitive to overconfident labels (Supplementary Section 6.1). For benchmark methods requiring discrete labels, we used the largest *y*_*ik*_ to assign each cell to one candidate type.

### 4.5 Model optimization

The encoder *E*, decoder *G*, classifier *C* and batch discriminator *D* were optimized with a learning rate of The classifier and discriminator were updated ten times for each update of the encoder and decoder. Each mini-batch contained 512 cells sampled across batches. Training used a 50-epoch warm-up period in which the adversarial batch loss and classifier loss were excluded. This allowed the VAE embedding to learn the main transcriptomic structure before batch-adversarial and label-guided objectives were introduced.

## Acknowledgements

This work is supported by NSF DMS2310836 and 5U01AI167892.

## Declarations

- Data availability: See Supplementary Material, Suppl Section 1
- Code availability: https://github.com/ch2343/scCRAFTplus

## References

[1] Qiu, C., Cao, J., Martin, B.K., Li, T., Welsh, I.C., Srivatsan, S., Huang, X., Calderon, D., Noble, W.S., Disteche, C.M., et al.: Systematic reconstruction of cellular trajectories across mouse embryogenesis. Nature genetics 54(3), 328–341 (2022)

[2] Travaglini, K.J., Nabhan, A.N., Penland, L., Sinha, R., Gillich, A., Sit, R.V., Chang, S., Conley, S.D., Mori, Y., Seita, J., et al.: A molecular cell atlas of the human lung from single-cell rna sequencing. Nature 587(7835), 619–625 (2020)

[3] Tritschler, S., Thomas, M., Böttcher, A., Ludwig, B., Schmid, J., Schubert, U., Kemter, E., Wolf, E., Lickert, H., Theis, F.J.: A transcriptional cross species map of pancreatic islet cells. Molecular Metabolism 66, 101595 (2022)

[4] Hao, Y., Hao, S., Andersen-Nissen, E., Mauck, W.M., Zheng, S., Butler, A., Lee, M.J., Wilk, A.J., Darby, C., Zager, M., et al.: Integrated analysis of multimodal single-cell data. Cell 184(13), 3573–3587 (2021)

[5] Lopez, R., Regier, J., Cole, M.B., Jordan, M.I., Yosef, N.: Deep generative modeling for single-cell transcriptomics. Nature methods 15(12), 1053–1058 (2018)

[6] Korsunsky, I., Millard, N., Fan, J., Slowikowski, K., Zhang, F., Wei, K., Baglaenko, Y., Brenner, M., Loh, P.-r., Raychaudhuri, S.: Fast, sensitive and accurate integration of single-cell data with harmony. Nature methods 16(12), 1289–1296 (2019)

[7] Hie, B., Bryson, B., Berger, B.: Efficient integration of heterogeneous single-cell transcriptomes using scanorama. Nature biotechnology 37(6), 685–691 (2019)

[8] He, C., Filippidis, P., Kleinstein, S.H., Guan, L.: Partially characterized topology guides reliable anchor-free scrna-integration. Communications Biology 8(1), 561 (2025)

[9] Ding, J., Adiconis, X., Simmons, S.K., Kowalczyk, M.S., Hession, C.C., Marjanovic, N.D., Hughes, T.K., Wadsworth, M.H., Burks, T., Nguyen, L.T., et al.: Systematic comparison of single-cell and single-nucleus rna-sequencing methods. Nature biotechnology 38(6), 737–746 (2020)

[10] Mereu, E., Lafzi, A., Moutinho, C., Ziegenhain, C., McCarthy, D.J., Álvarez-Varela, A., Batlle, E., Sagar, n., Gruen, D., Lau, J.K., et al.: Benchmarking single-cell rna-sequencing protocols for cell atlas projects. Nature biotechnology 38(6), 747–755 (2020)

[11] Hrovatin, K., Sikkema, L., Shitov, V.A., Heimberg, G., Shulman, M., Oliver, A.J., Mueller, M.F., Ibarra, I.L., Wang, H., Ramírez-Suástegui, C., et al.: Considerations for building and using integrated single-cell atlases. Nature Methods, 1–17 (2024)

[12] Andreatta, M., Berenstein, A.J., Carmona, S.J.: scgate: marker-based purification of cell types from heterogeneous single-cell rna-seq datasets. Bioinformatics 38(9), 2642–2644 (2022)

[13] Andreatta, M., Hérault, L., Gueguen, P., Gfeller, D., Berenstein, A.J., Carmona, S.J.: Semi-supervised integration of single-cell transcriptomics data. Nature Communications 15(1), 872 (2024)

[14] Lin, Y., Cao, Y., Kim, H.J., Salim, A., Speed, T.P., Lin, D.M., Yang, P., Yang, J.Y.H.: scclassify: sample size estimation and multiscale classification of cells using single and multiple reference. Molecular systems biology 16(6), 9389 (2020)

[15] Rood, J.E., Maartens, A., Hupalowska, A., Teichmann, S.A., Regev, A.: Impact of the human cell atlas on medicine. Nature medicine 28(12), 2486–2496 (2022)

[16] Hrovatin, K., Bastidas-Ponce, A., Bakhti, M., Zappia, L., Büttner, M., Salinno, C., Sterr, M., Böttcher, A., Migliorini, A., Lickert, H., et al.: Delineating mouse β-cell identity during lifetime and in diabetes with a single cell atlas. Nature metabolism 5(9), 1615–1637 (2023)

[17] Novella-Rausell, C., Grudniewska, M., Peters, D.J., Mahfouz, A.: A comprehensive mouse kidney atlas enables rare cell population characterization and robust marker discovery. Iscience 26(6) (2023)

[18] Sikkema, L., Ramírez-Suástegui, C., Strobl, D.C., Gillett, T.E., Zappia, L., Madissoon, E., Markov, N.S., Zaragosi, L.-E., Ji, Y., Ansari, M., et al.: An integrated cell atlas of the lung in health and disease. Nature medicine 29(6), 1563–1577 (2023)

[19] Peterson, V.M., Zhang, K.X., Kumar, N., Wong, J., Li, L., Wilson, D.C., Moore, R., McClanahan, T.K., Sadekova, S., Klappenbach, J.A.: Multiplexed quantification of proteins and transcripts in single cells. Nature biotechnology 35(10), 936–939 (2017)

[20] Efremova, M., Teichmann, S.A.: Computational methods for single-cell omics across modalities. Nature methods 17(1), 14–17 (2020)

[21] Stoeckius, M., Hafemeister, C., Stephenson, W., Houck-Loomis, B., Chattopadhyay, P.K., Swerdlow, H., Satija, R., Smibert, P.: Simultaneous epitope and transcriptome measurement in single cells. Nature methods 14(9), 865–868 (2017)

[22] Zhang, A.W., O’Flanagan, C., Chavez, E.A., Lim, J.L., Ceglia, N., McPherson, A., Wiens, M., Walters, P., Chan, T., Hewitson, B., et al.: Probabilistic cell-type assignment of single-cell rna-seq for tumor microenvironment profiling. Nature methods 16(10), 1007–1015 (2019)

[23] Zhang, Z., Luo, D., Zhong, X., Choi, J.H., Ma, Y., Wang, S., Mahrt, E., Guo, W., Stawiski, E.W., Modrusan, Z., et al.: Scina: a semi-supervised subtyping algorithm of single cells and bulk samples. Genes 10(7), 531 (2019)

[24] Andreatta, M., Carmona, S.J.: Ucell: Robust and scalable single-cell gene signature scoring. Computational and structural biotechnology journal 19, 3796–3798 (2021)

[25] Abdelaal, T., Michielsen, L., Cats, D., Hoogduin, D., Mei, H., Reinders, M.J., Mahfouz, A.: A comparison of automatic cell identification methods for single-cell rna sequencing data. Genome biology 20, 1–19 (2019)

[26] Xu, C., Lopez, R., Mehlman, E., Regier, J., Jordan, M.I., Yosef, N.: Probabilistic harmonization and annotation of single-cell transcriptomics data with deep generative models. Molecular systems biology 17(1), 9620 (2021)

[27] Hu, J., Li, X., Hu, G., Lyu, Y., Susztak, K., Li, M.: Iterative transfer learning with neural network for clustering and cell type classification in single-cell rna-seq analysis. Nature machine intelligence 2(10), 607–618 (2020)

[28] Shree, A., Pavan, M.K., Zafar, H.: scdreamer for atlas-level integration of single-cell datasets using deep generative model paired with adversarial classifier. Nature Communications 14(1), 7781 (2023)

[29] Lotfollahi, M., Wolf, F.A., Theis, F.J.: scgen predicts single-cell perturbation responses. Nature methods 16(8), 715–721 (2019)

[30] Franzén, O., Gan, L.-M., Björkegren, J.L.: Panglaodb: a web server for exploration of mouse and human single-cell rna sequencing data. Database 2019, 046 (2019)

[31] Zhang, X., Lan, Y., Xu, J., Quan, F., Zhao, E., Deng, C., Luo, T., Xu, L., Liao, G., Yan, M., et al.: Cellmarker: a manually curated resource of cell markers in human and mouse. Nucleic acids research 47(D1), 721–728 (2019)

[32] Pasquini, G., Arias, J.E.R., Schäfer, P., Busskamp, V.: Automated methods for cell type annotation on scrna-seq data. Computational and Structural Biotechnology Journal 19, 961–969 (2021)

[33] Luecken, M.D., Theis, F.J.: Current best practices in single-cell rna-seq analysis: a tutorial. Molecular systems biology 15(6), 8746 (2019)

[34] Shen, X., He, C., Guan, L.: A benchmark of semi-supervised scrna-seq integration methods in real-world scenarios. PLOS Computational Biology 22(3), 1014008 (2026)

[35] Miyato, T., Maeda, S.-i., Koyama, M., Ishii, S.: Virtual adversarial training: a regularization method for supervised and semi-supervised learning. IEEE transactions on pattern analysis and machine intelligence 41(8), 1979–1993 (2018)

[36] Wolf, F.A., Angerer, P., Theis, F.J.: Scanpy: large-scale single-cell gene expression data analysis. Genome biology 19, 1–5 (2018)

