## Supplementary file for "Robust semi-supervised scRNA-seq integration from virtual adversarial learning"

**6 Cell type prediction with wrong marker list** **15**

**7 Computational Cost, Efficiency, and Scaling with Cell and Marker Count** **16**

### 1 Dataset details

In our study, we applied scCRAFT+ to integrate four diverse real datasets, presenting performance over various tissues (Table S1). These datasets included two PBMC dataset (PBMC and human immune), one pancreatic dataset (Pancreatic Islets) and one lung dataset (Lung Atlas). The ground truth for these datasets was based on original annotations from the source datasets, consistent with other benchmarking studies.

**Table S1** Integration Task Summary

| Integration Task | Cell Number | Batches | Reference Tissue |
|---|---|---|---|
| Human PBMC | 26128 | 8 | Human - PBMC |
| Human immune | 33506 | 10 | Human - PBMC |
| Lung | 29256 | 16 | Human - Lung v2 (HLCA) |
| Pancreatic Islet | 16382 | 9 | Human - Pancreas |

Note: This table summarizes the integration tasks highlighting cell numbers, batches and reference tissue used for the auto-annotation together with integration.

#### Human PBMC

The PBMC dataset includes cells sequenced from two distinct experiments and five single-cell RNA sequencing protocols, sourced from Ding et al. [1] and available through the Single Cell Portal at the Broad Institute https://singlecell.broadinstitute.org/single_cell/study/SCP424/single-cell-comparison-pbmc-data. We primarily analyzed data from the second experiment, as it provides a more balanced sample size across four batches, allowing us to focus on single-level batch effects to enhance imputation precision. After quality control filtering, the dataset consists of 26,128 cells and 22,891 genes across the selected batches.

##### Human Immune dataset

The Human Immune dataset includes cells derived from ten human samples across two distinct tissues: bone marrow and peripheral blood. This dataset encompasses a range of experimental protocols and sampling strategies designed to capture the diversity of immune cell types within these tissues. Comprehensive details on dataset procurement, the varied experimental protocols employed, and the criteria for sample selection are provided by Luecken et al. [2]. Following rigorous quality control measures, the dataset was refined to include 33,506 cells across 12,303 genes, spanning ten different batches.

##### Pancreatic Islet dataset

The Pancreatic Islet dataset incorporates five publicly available pancreatic islet datasets, creating a comprehensive collection of 16,382 cells aggregated from multiple studies [3–7]. Detailed documentation on the compilation process, including the criteria for cell selection, quality control measures, and experimental protocols for each batch, is provided by Luecken et al. [2], ensuring transparency and reproducibility in our analysis.

##### Lung atlas dataset

The Lung Atlas dataset, sourced from Vieira Braga et al.[8] and archived in the Gene Expression Omnibus (GEO) under accession code GSE130148, offers a detailed exploration of the cellular landscape within lung tissue. Following stringent quality control, the dataset includes 29,256 cells and 15,139 genes, providing a robust foundation for lung cell-type characterization. Comprehensive documentation on dataset assembly, encompassing quality control protocols, experimental methodologies applied across batches, and acquisition specifics, is provided by Luecken et al.[2], ensuring rigorous standards of transparency and reproducibility for integrative analysis and cross-study comparisons in lung research.

#### 2 Benchmarking Methods

In our benchmarking study, we evaluated three semi-supervised computational methods for single-cell data integration, focusing on their ability to leverage the cell type information while be robust to the noisy annotation. These methods can be categorized into two main approaches: deep neural network frameworks with cell type information-guided embedding generation and cell type-informed nearest neighbor filtering.

Deep neural network frameworks with cell type information-guided embedding generation, such as scGen [9] and scANVI [10], utilize neural networks to generate embeddings that integrate cell type information in different ways. scGen implicitly learns cell type differences through a variational autoencoder that captures cell type and batch variations without directly using cell type labels in the loss function. In contrast, scANVI explicitly incorporates cell type information by adding a classification loss, allowing the model to use partial cell type annotations for enhanced embedding generation, particularly in semi-supervised settings.

Cell type-informed nearest neighbor filtering, exemplified by ssSTACAS [11], builds on traditional integration frameworks but adds a filtering step based on cell type annotations. By retaining only nearest neighbor links within the same cell type, ssSTACAS achieves a biologically consistent alignment, reducing batch effects while preserving clear cell type boundaries in the integrated data.

A summary of the methods used, including version information, specific settings, and tutorial references, is provided in Table S2. We also include Harmony [12], seurat [13], scVI [14] and scCRAFT [15] as the unsupervised benchmarking competitors with detailed specification and tutorial details in He. et al [15].

**Table S2** Summary of Benchmarking Methods

| Method | Specific Settings | Tutorial Website |
|---|---|---|
| scANVI | 50-dimensional latent space, 128 nodes per hidden layer, 2 layers with maximum 50 training epochs | https://docs.scvi-tools.org/en/stable/tutorials/notebooks/scrna/seed_labeling.html |
| scGen | maximum 50 training epochs | https://scgen.readthedocs.io/en/stable/tutorials/scgen_batch_removal.html |
| ssSTACAS | top 50 PCA embeddings | https://github.com/carmonalab/STACAS |

* Note: All methods were configured to output a 50-dimensional embedding.

### 3 Evaluation Metrics

#### 3.1 Overview

To evaluate single-cell data integration methods, we focused on two key areas: (1) preservation of biological variation and (2) batch effect removal. Metrics for biological conservation assess the extent to which each method retains the underlying biological signals, while batch correction metrics measure effectiveness in reducing batch-induced artifacts. To obtain a comprehensive assessment, composite scores were calculated as the weighted average of metrics within each category. The Overall Score combines these two composite scores, with weights reflecting their relative importance in single-cell integration. Further details on each metric and calculation are provided in He et al. [15].

Table S3 provides an overview of the key metrics used in our evaluation, including the composite scores that combine these metrics into a single value. To emphasize their importance in the overall assessment, composite scores are highlighted in bold.

**Table S3** Summary of Evaluation Metrics

| Metric | Description |
|---|---|
| **Biological Variance Conservation** | |
| Normalized Mutual Information (NMI) | Measures overlap between cell-type labels and clusters. |
| Adjusted Rand Index (ARI) | Assesses clustering accuracy relative to known labels. |
| Cell Type Average Silhouette Width (ASW_cell) | Evaluates cluster cohesion and separation. |
| **Batch Effect Correction** | |
| Average Silhouette Width for Batches (ASW_batch) | Measures distance between different batches. |
| Graph Connectivity | Evaluates cohesiveness of cells of the same type within the kNN graph. |
| k-Nearest Neighbor Batch Effect Test (kBET) | Assesses local neighborhood consistency with global label distribution. |
| Batch Label Identity Score Index (bLISI) | Measures batch diversity within local neighborhoods. |
| Proportion of True Positive Cells | Assesses the proportion of positive cells which mirror the global batch distribution. |
| **Composite Scores** | |
| **Biological Conservation Score** | Average of NMI, ARI and ASW_cell. |
| **Batch Mixing Score** | Average of bLISI, ASW_batch, kBET, Graph Connectivity, and true positive rate. |
| **Overall Score** | Weighted sum of Biological Conservation Score (60%) and Batch Mixing Score (40%). |

### 4 UMAP evaluation for four real datasets

The arrangement of panels is the same across Fig. SS1-S4, and we provide their shared Figure legends here to avoid redundancy:

UMAP visualization contrasting the unintegrated dataset with its integrated counterparts using all semi-supervised and unsupervised methods.

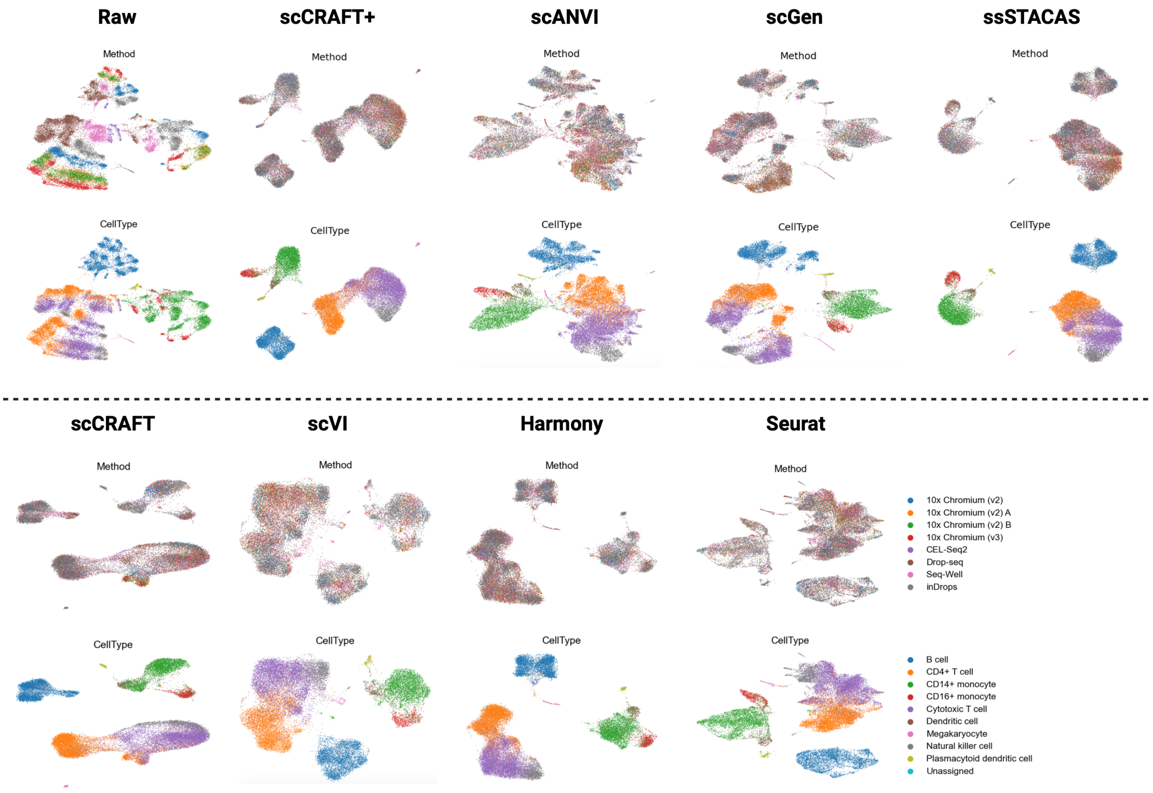

**Fig. S1** PBMC Dataset Results.

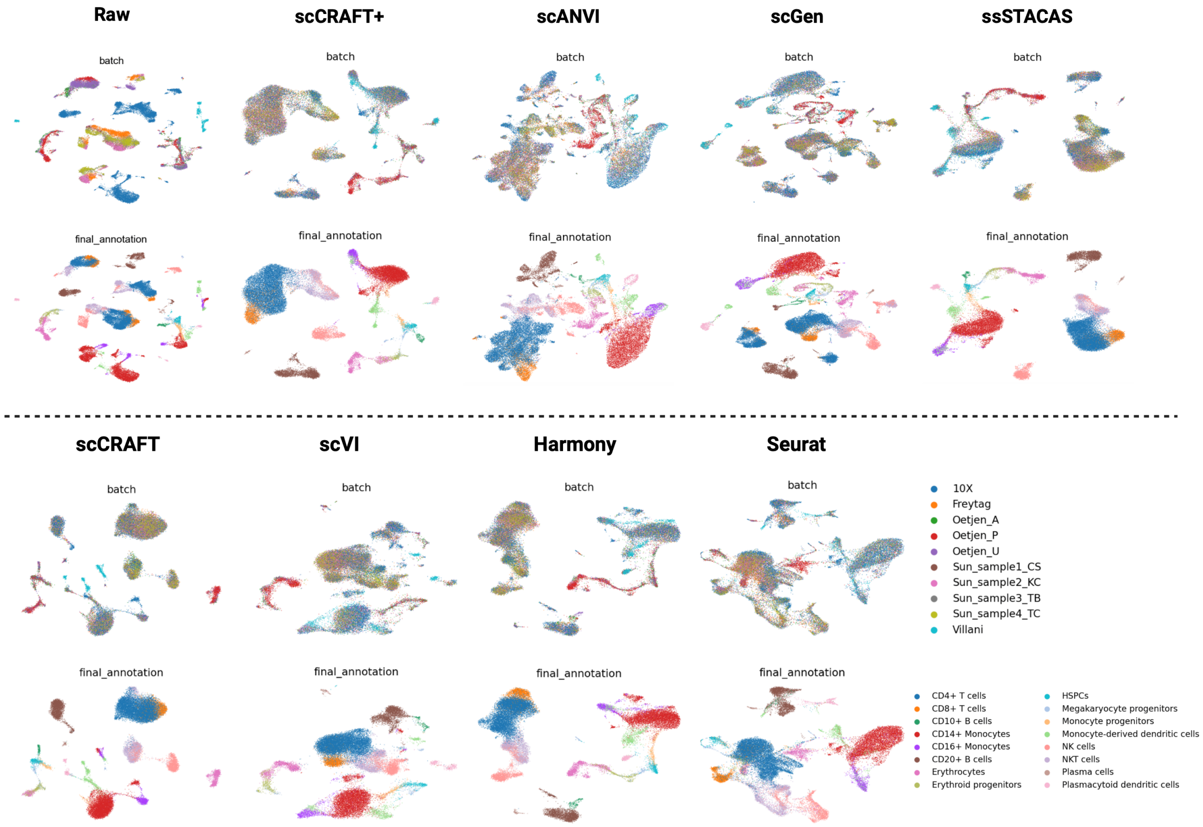

**Fig. S2** Human Immune Dataset Results.

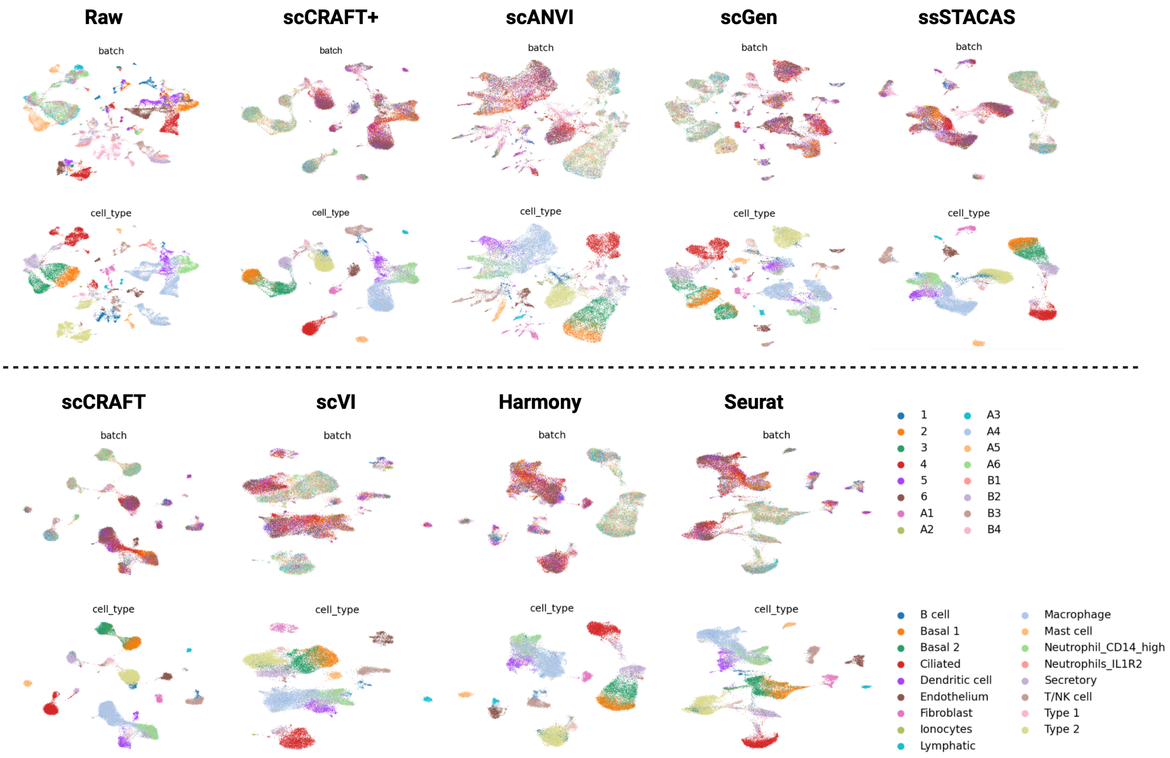

**Fig. S3** Lung Atlas Dataset Results

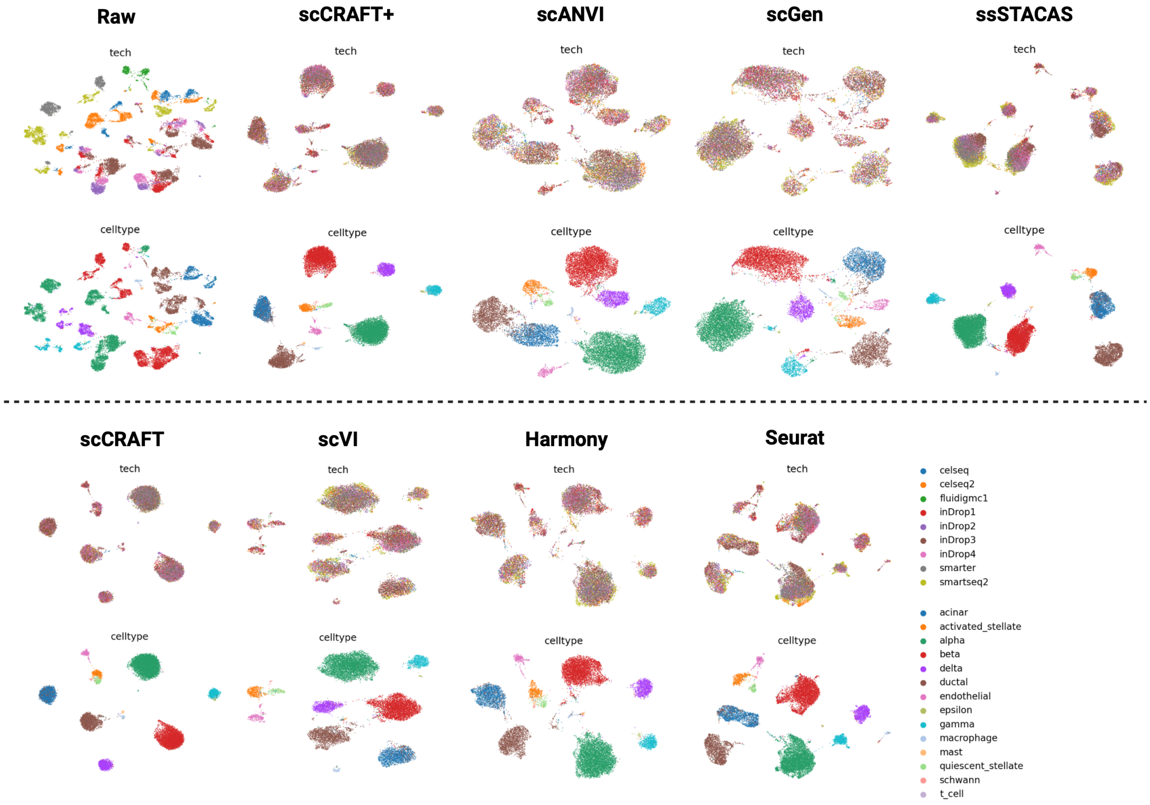

**Fig. S4** Pancreas Dataset Results.

### 5 Ablation Study

#### 5.1 Robustness of VAT to Label Assignment Confidence

We investigate how VAT contributes to scCRAFT+'s robustness under varying degrees of label assignment confidence. In scCRAFT+, soft labels are constructed from U scores, which quantify graded cell-type affinities based on marker gene expression. By tuning the temperature parameter $t$ in the sharpening operation, we can adjust the label certainty: lower $t$ yields more confident (harder) labels, while higher $t$ retains more ambiguity.

To systematically assess the role of VAT under different label sharpness conditions, we benchmark integration performance under four scenarios:

1. $t = 5$: nearly plain cell type information (very soft labels);
2. $t = 1$: unsharpened soft labels (low confidence);
3. $t = 0.05$: default training value (moderate-to-high confidence);
4. $t = 0$: near one-hot hard labels.

We compare model performance with and without VAT under each of these conditions using biological conservation, batch correction, and composite scores. The results are shown in Table S4.

**Table S4** Comparison of VAT vs. No-VAT under Varying Label Assignment Confidence ($t$)

| | Method | PBMC | Human Immune | Lung Atlas | Pancreas |
|---|---|---|---|---|---|
| | | **Biological Conservation** | | | |
| $t = 5$ | No VAT | 0.6452 | 0.7154 | 0.6924 | 0.8579 |
| | VAT | 0.6526 | 0.7206 | 0.6916 | 0.8542 |
| $t = 1$ | No VAT | 0.6658 | 0.7274 | 0.6946 | 0.8541 |
| | VAT | 0.6686 | 0.7238 | 0.6928 | 0.8552 |
| $t = 0.05$ | No VAT | 0.6961 | 0.7337 | 0.7041 | 0.8572 |
| | VAT | 0.7036 | 0.7506 | 0.6964 | 0.8598 |
| $t = 0$ | No VAT | 0.5822 | 0.6052 | 0.5934 | 0.6933 |
| | VAT | 0.6689 | 0.7474 | 0.6986 | 0.8577 |
| | | **Batch Correction** | | | |
| $t = 5$ | No VAT | 0.5618 | 0.6013 | 0.5742 | 0.7256 |
| | VAT | 0.5924 | 0.6283 | 0.6154 | 0.7312 |
| $t = 1$ | No VAT | 0.5801 | 0.6026 | 0.5689 | 0.7240 |
| | VAT | 0.6294 | 0.6491 | 0.6265 | 0.7396 |
| $t = 0.05$ | No VAT | 0.6230 | 0.6410 | 0.6010 | 0.7170 |
| | VAT | 0.6315 | 0.6171 | 0.5995 | 0.7448 |
| $t = 0$ | No VAT | 0.6145 | 0.6020 | 0.5880 | 0.6890 |
| | VAT | 0.6158 | 0.6130 | 0.5984 | 0.6923 |
| | | **Overall Scores** | | | |
| $t = 5$ | No VAT | 0.6118 | 0.6698 | 0.6451 | 0.7858 |
| | VAT | 0.6295 | 0.6837 | 0.6619 | 0.8050 |
| $t = 1$ | No VAT | 0.6315 | 0.6779 | 0.6424 | 0.8088 |
| | VAT | 0.6529 | 0.6939 | 0.6663 | 0.8083 |
| $t = 0.05$ | No VAT | 0.6677 | 0.6966 | 0.6623 | 0.8011 |
| | VAT | 0.6748 | 0.6972 | 0.6576 | 0.8138 |
| $t = 0$ | No VAT | 0.5831 | 0.6043 | 0.5912 | 0.6908 |
| | VAT | 0.6477 | 0.6937 | 0.6585 | 0.7916 |

As shown in Table S4, the effectiveness of scCRAFT+ under different sharpening temperatures highlights the critical role of VAT in handling varying degrees of label confidence. When $t = 0$, corresponding to hard label assignments, models trained without VAT tend to overfit to discrete label boundaries, leading to oversegregated embeddings and fragmented biological structures. In contrast, VAT helps mitigate this rigidity by enforcing local consistency, resulting in improved biological conservation and overall integration performance.

On the other hand, when $t = 1$, label assignments become overly diffuse, and the supervision signal effectively fades. In this case, both VAT and no-VAT models converge toward the behavior of the unsupervised scCRAFT baseline, as the training no longer benefits meaningfully from label guidance. When the labels are softened even further ($t = 5$), the supervision becomes nearly uninformative, and the model again approaches the behavior of plain scCRAFT without marker information. Nonetheless, VAT still provides a modest gain compared with the no-VAT counterpart by regularizing the latent space. These performance differences are evident in human immune and PBMC datasets, where the integration quality benefits more from the assigned label.

For completeness, at the intermediate setting $t = 0.05$, scCRAFT+ with VAT achieves optimal integration by balancing label certainty with flexibility, but the clearest insights into VAT's robustness emerge from its ability to stabilize performance across both low-certainty ($t = 1$ or $t = 5$) and high-certainty ($t = 0$) regimes.

To visualize the effect of label sharpening, we examine the U score distributions for $NK_2$ cells across three $t$ values in Figure S5. As $t$ decreases, U scores become increasingly peaked, transitioning from soft probabilistic assignments to near one-hot encodings. Despite these changes, VAT ensures consistent integration performance by regularizing local structure in the embedding space.

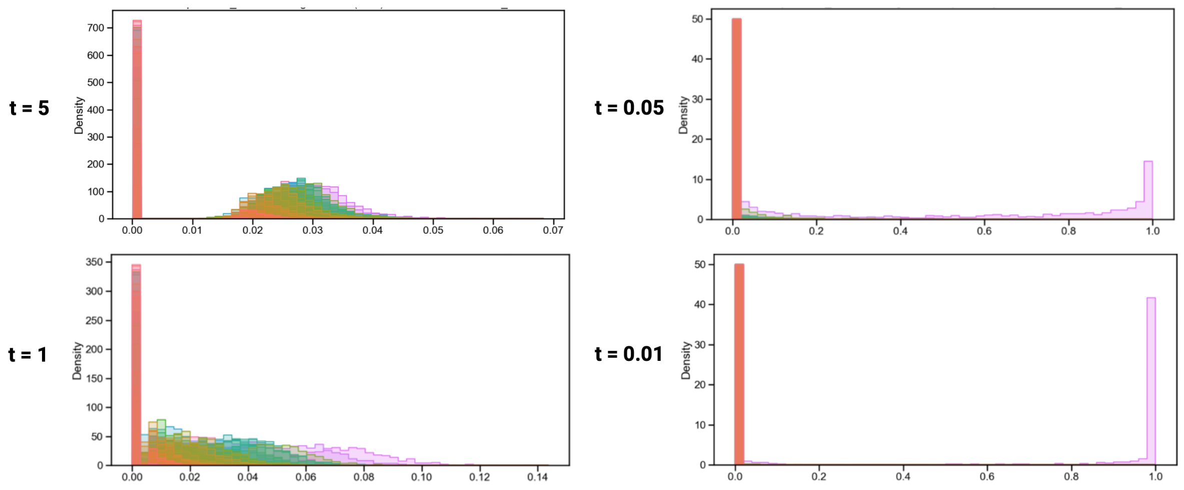

**Fig. S5** Effect of temperature sharpening on U scores for $NK_2$ cells. Each panel shows the distribution of sharpened U scores across cell types for a different temperature parameter: (upper left) $t = 5$, (lower left) $t = 1$, (upper right) $t = 0.05$, and (lower right) $t = 0.01$. Lower $t$ values produce sharper, more confident label assignments.

In summary, these results highlight the flexibility of U score–based labeling and confirm that scCRAFT+ equipped with VAT is robust to varying levels of label confidence. This enables users to apply the method without needing to fine-tune the sharpening temperature for each new dataset. In contrast, models trained without VAT are more brittle: performance declines when label confidence is too high or too low, underscoring the importance of VAT in handling uncertain or noisy annotations in single-cell integration tasks.

#### 5.2 VAT improves cluster-level consistency

We first examined the effect of VAT under hard-label supervision. Without VAT, the model tended to produce over-fragmented predicted labels, splitting local regions of the embedding into small subclusters that were not well supported by the broader transcriptomic structure (Fig. S6a). In contrast, VAT reduced these fragmented predictions by enforcing local prediction consistency, leading to annotations that better followed the major embedding structure.

Under soft-label supervision, both VAT and no-VAT models showed less severe fragmentation than in the hard-label setting, reflecting the benefit of using marker-derived probabilistic labels rather than fixed one-hot assignments (Fig. S6b). However, visual inspection alone does not fully capture whether the predicted labels are well aligned with the embedding-derived cluster structure. We therefore performed an additional cluster-level consistency analysis under the soft-label setting.

This analysis is distinct from the integration evaluation in Table S4. In Table S4, ARI and NMI are used as biological-conservation metrics by comparing embedding-derived clusters with the source-provided cell type labels. Here, ARI and NMI instead evaluate prediction coherence by comparing embedding-derived clusters with the model-predicted labels. For each dataset and model variant, we performed unsupervised clustering over a range of resolutions and reported the best ARI and NMI between the resulting clusters and the predicted labels. Under this analysis, VAT improved the optimal ARI by an average of 14.1% and the optimal NMI by an average of 18.9% compared with the no-VAT model, indicating that VAT-guided predictions are more consistent with the transcriptomic cluster structure.

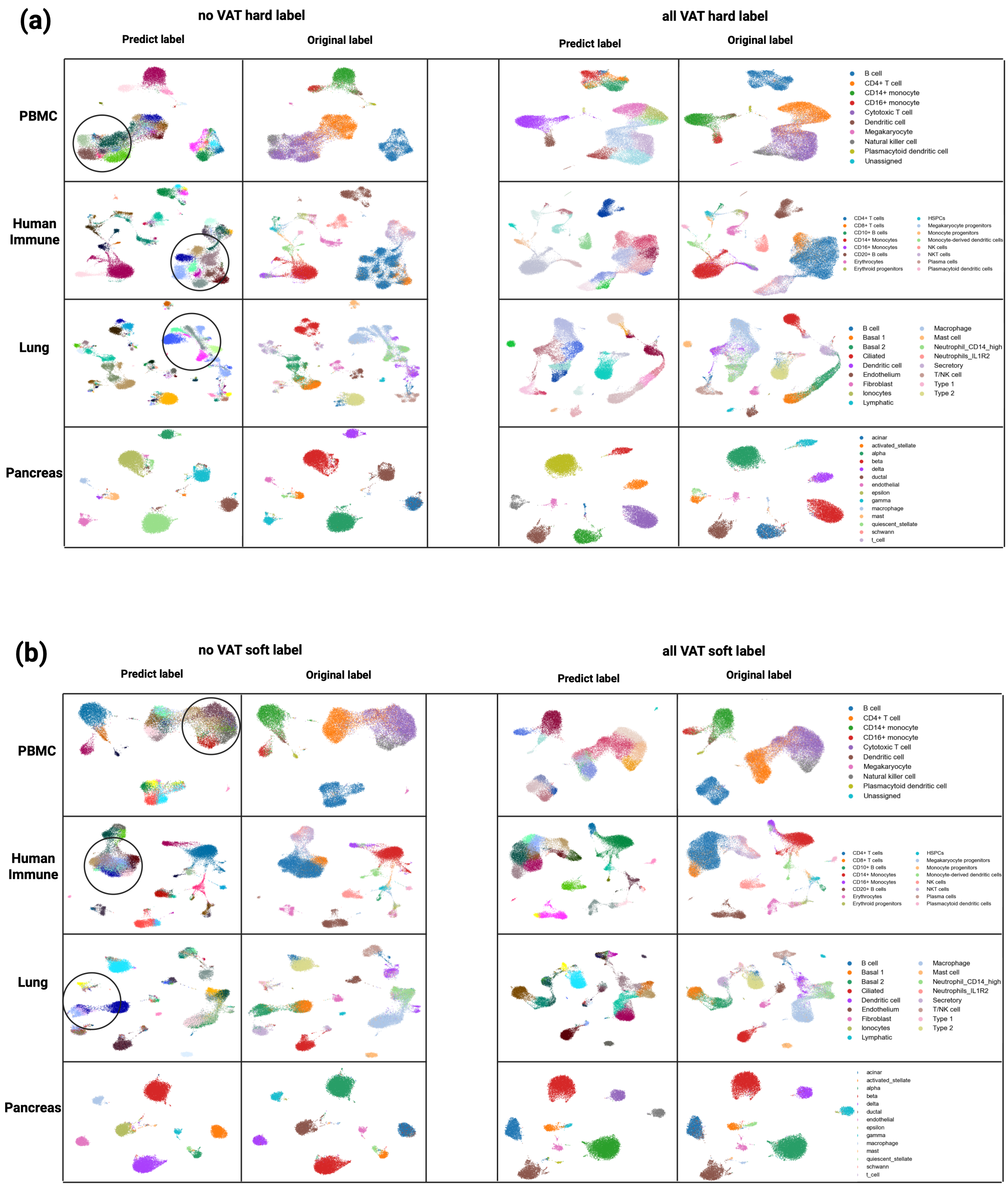

**Fig. S6** UMAP visualizations of scCRAFT+ integration results across four datasets. Cells are colored by model-predicted labels and source-provided reference labels. (a) Comparison under hard-label supervision, where VAT reduces over-fragmented predicted labels and improves consistency with the embedding structure. (b) Comparison under soft-label supervision, where VAT further improves the alignment between predicted labels and embedding-derived clusters.

#### 5.3 VAT ablation study within scVI framework

We extended our VAT framework to other deep learning models, including scVI. Given that scVI's semi-supervised variant, scANVI, employs cross-entropy loss to leverage cell type labels, it provides a suitable benchmark for comparing cross-entropy with the VAT framework. In this study, we implemented a VAT-based version of scANVI, replacing its cross-entropy loss with the VAT framework. Using auto-annotated hard labels from the Fast U score (Section 2.3) as reference, we benchmarked scANVI+VAT against the original scANVI. As shown in Fig. S2, the VAT-enhanced model achieved comparable batch correction while demonstrating significant improvements in biological conservation, with gains of +4.0% (PBMC), +3.6% (human immune), +3.6% (lung atlas), and +2.1% (pancreas) across datasets. Notably, these improvements approached but did not exceed scCRAFT+'s performance (Fig. S2), confirming VAT's ability to enhance biological resolution without compromising integration quality.

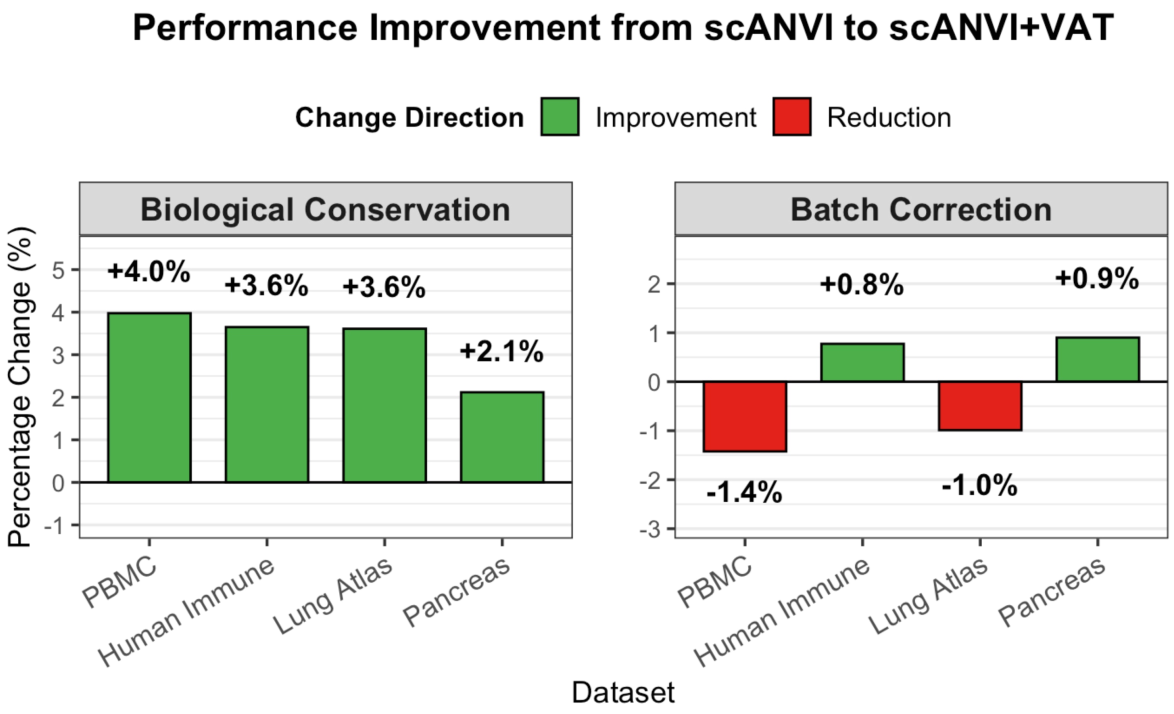

**Fig. S7** Performance comparison of scANVI and scANVI+VAT. Bar plots contrast (left) biological conservation and (right) batch correction scores across four datasets.

#### 5.4 Sensitivity analysis for the VAT perturbation radius

In the VAT objective, the hyperparameters $\alpha$ and $\epsilon$ control the strength of local prediction smoothing. In this study, we fixed the LDS weight at $\alpha = 1$ and focused on selecting the perturbation radius $\epsilon$, which determines how far the classifier is perturbed along the most prediction-sensitive direction in the latent embedding space. Intuitively, a smaller $\epsilon$ imposes weaker local smoothness and may retain fragmented, mosaic-like predictions, whereas an overly large $\epsilon$ may over-smooth the classifier and merge smaller cell populations.

To examine this trade-off, we evaluated $\epsilon \in \{10, 50, 100, 500, 1000\}$ across four datasets. For each setting, we quantified prediction smoothness by the mean nearest-neighbor label agreement in the learned embedding and measured annotation granularity by the number of predicted labels (Fig. S8). Increasing $\epsilon$ generally improved local prediction smoothness, and $\epsilon = 500$ achieved the peak smoothness in three out of four datasets or reached a near-maximum level. At the same time, $\epsilon = 500$ substantially reduced the number of predicted labels compared with $\epsilon = 100$, indicating that many fragmented or mosaic-like assignments were smoothed into coherent cluster-level annotations. Further increasing $\epsilon$ to 1000 did not consistently improve smoothness and sometimes reduced annotation diversity, suggesting the onset of over-smoothing. Therefore, we selected $\epsilon = 500$ as a practical balance between improving local prediction coherence and preserving biologically meaningful subtype structure.

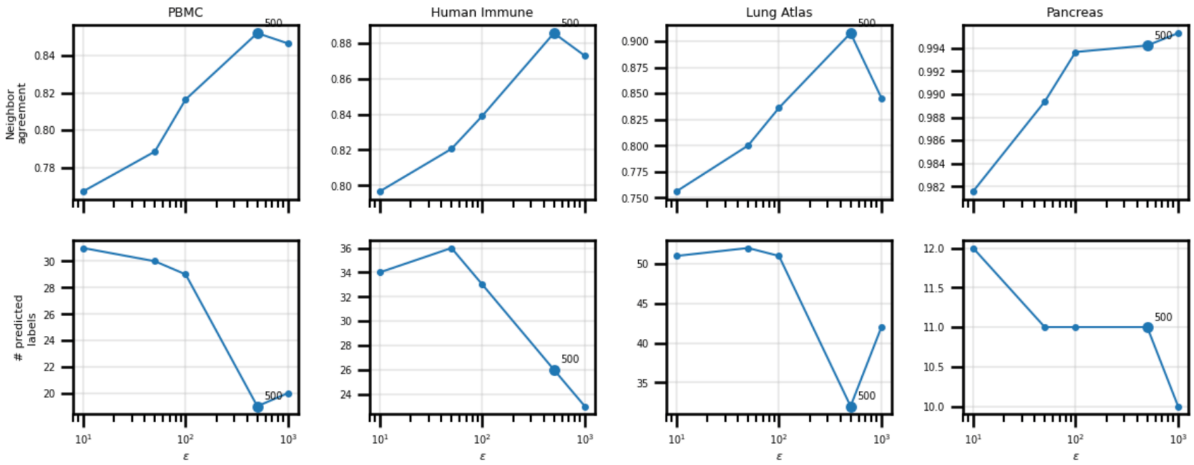

**Fig. S8** Sensitivity analysis of the VAT perturbation radius $\epsilon$. The top row shows mean nearest-neighbor label agreement, which measures local prediction smoothness in the learned embedding. The bottom row shows the number of predicted labels, reflecting annotation granularity. Across datasets, increasing $\epsilon$ improves prediction smoothness, while larger values reduce fragmented predicted labels. $\epsilon = 500$ provides a balance between local annotation coherence and preservation of subtype diversity.

#### 6 Cell type prediction with wrong marker list

We further examined how scCRAFT+ behaves when the marker reference is not tissue-matched. The goal of this analysis is not to claim that mismatched markers provide definitive annotations, but to evaluate whether scCRAFT+ can still extract useful partial information while avoiding over-confident misclassification.

When lung markers were applied to the human immune dataset, scCRAFT+ recovered broad immune compartments, including B cell, T cell, dendritic cell, and monocyte-like populations (Fig. S9a). Although these annotations are less detailed than tissue-matched immune annotations, they provide a useful first-pass classification of major shared cell types. This suggests that even an imperfect marker list can still offer informative coarse-grained guidance when it contains markers for cell types present in the target data.

In contrast, when PBMC markers were applied to the lung atlas dataset, only immune-related populations were confidently annotated, while many lung-resident non-immune populations were assigned as "Unknown" (Fig. S9b). This behavior is expected because PBMC markers do not contain appropriate references for many lung-specific cell types, such as epithelial, endothelial, or fibroblast populations. Consistently, the proportion of "Unknown" predictions increased to 74.4% when PBMC markers were used for lung annotation, whereas it was only 0.3% when the matched lung marker list was used. Thus, a high fraction of "Unknown" predictions can provide users with a practical warning that the supplied marker reference may be poorly matched to the target tissue.

Together, these examples illustrate two practical outcomes of using mismatched marker lists: scCRAFT+ can provide coarse annotation for shared cell populations, and it can flag cell populations not supported by the reference marker list as uncertain rather than assigning over-confident incorrect labels.

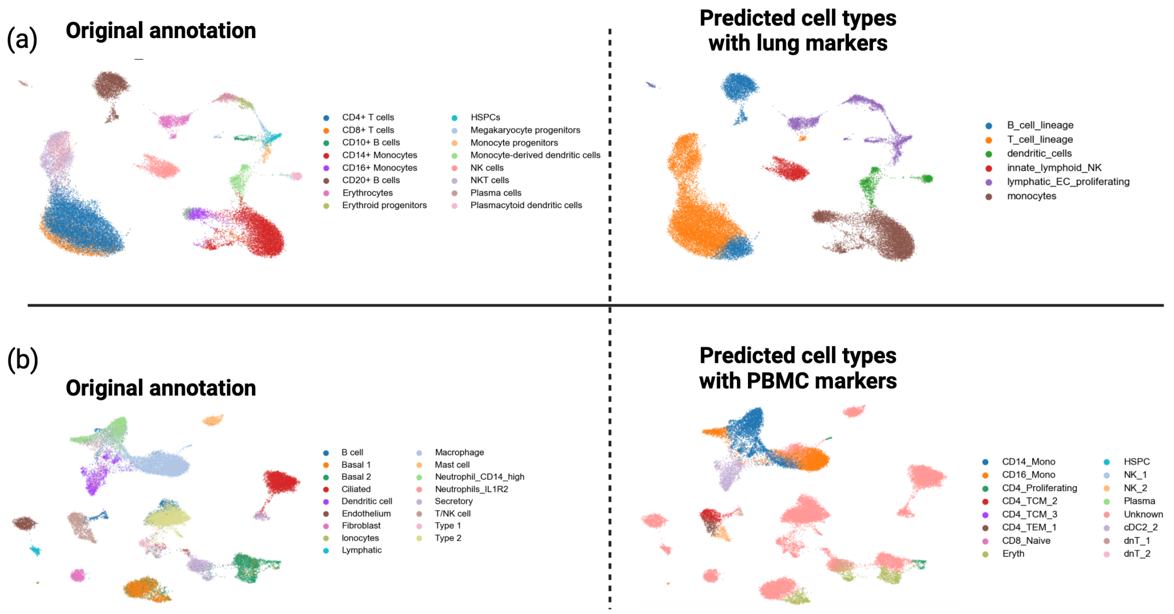

**Fig. S9** Wrong marker list auto-annotation. (a) UMAP visualization of scCRAFT+ integration embeddings of the human immune dataset, colored by original annotation and predicted cell types using lung markers. Lung markers provide coarse annotation of shared immune compartments. (b) UMAP visualization of scCRAFT+ integration embeddings of the lung atlas dataset, colored by original annotation and predicted cell types using PBMC markers. PBMC markers annotate immune-related populations while leaving unsupported lung-resident populations as "Unknown".

### 7 Computational Cost, Efficiency, and Scaling with Cell and Marker Count

The marker-based preprocessing in scCRAFT+ consists of two core steps: (i) computing rank statistics for marker genes, and (ii) aggregating these ranks into per-cell U scores for each candidate cell type. Below we summarize the computational cost and scaling behavior with increasing numbers of cells and markers.

#### Precomputing ranks for marker genes

**Implementation note.** In our implementation, we construct a matrix of expression values restricted to the union of marker genes across all cell types, and compute gene-wise ranks across cells using `pandas.DataFrame.rank()` (default `axis=0`). Therefore, ranks are computed *across cells for each marker gene*, rather than across genes within each cell.

**Computational complexity.** Let $C$ be the number of cells and $M$ the number of unique marker genes (after intersecting with the dataset). Ranking each marker gene across $C$ cells has complexity $O(C \log C)$, yielding an overall cost of

$$O(M\, C \log C).$$

Memory usage is $O(CM)$ to store the marker expression and ranks.

**Scaling behavior.** The dominant factor is the cell count $C$, with an additional logarithmic factor from ranking. Increasing the marker pool size $M$ increases both runtime and memory approximately linearly.

#### U-score computation and hard-label assignment

**Computational complexity.** Given precomputed ranks, U-score computation for each cell type requires summing over its marker list. Across all cell types, the total cost is linear in the number of cells and the number of marker entries used:

$$O(C \times M),$$

up to constant factors determined by overlap of marker sets. Selecting the maximum-scoring cell type per cell to obtain hard labels adds an additional $O(C \times K)$ operation where $K$ is the number of candidate cell types, typically negligible relative to rank computation.

**Scaling behavior.** This step scales approximately linearly with the number of cells and the total marker entries used, and is substantially faster than the rank-precomputation step.

#### Practical runtime

The marker preprocessing step is lightweight in practice. For a dataset with 100,000 cells using PBMC marker sets on an Intel 32-core CPU, the end-to-end U-score preprocessing completes in under 5 seconds.

### References

[1] Ding, J., Adiconis, X., Simmons, S.K., Kowalczyk, M.S., Hession, C.C., Marjanovic, N.D., Hughes, T.K., Wadsworth, M.H., Burks, T., Nguyen, L.T., *et al.*: Systematic comparison of single-cell and single-nucleus rna-sequencing methods. Nature biotechnology **38**(6), 737–746 (2020)

[2] Luecken, M.D., Büttner, M., Chaichoompu, K., Danese, A., Interlandi, M., Müller, M.F., Strobl, D.C., Zappia, L., Dugas, M., Colomé-Tatché, M., *et al.*: Benchmarking atlas-level data integration in single-cell genomics. Nature methods **19**(1), 41–50 (2022)

[3] Baron, M., Veres, A., Wolock, S.L., Faust, A.L., Gaujoux, R., Vetere, A., Ryu, J.H., Wagner, B.K., Shen-Orr, S.S., Klein, A.M., *et al.*: A single-cell transcriptomic map of the human and mouse pancreas reveals inter-and intra-cell population structure. Cell systems **3**(4), 346–360 (2016)

[4] Muraro, M.J., Dharmadhikari, G., Grün, D., Groen, N., Dielen, T., Jansen, E., Van Gurp, L., Engelse, M.A., Carlotti, F., De Koning, E.J., *et al.*: A single-cell transcriptome atlas of the human pancreas. Cell systems **3**(4), 385–394 (2016)

[5] Segerstolpe, Å., Palasantza, A., Eliasson, P., Andersson, E.-M., Andréasson, A.-C., Sun, X., Picelli, S., Sabirsh, A., Clausen, M., Bjursell, M.K., *et al.*: Single-cell transcriptome profiling of human pancreatic islets in health and type 2 diabetes. Cell metabolism **24**(4), 593–607 (2016)

[6] Lawlor, N., George, J., Bolisetty, M., Kursawe, R., Sun, L., Sivakamasundari, V., Kycia, I., Robson, P., Stitzel, M.L.: Single-cell transcriptomes identify human islet cell signatures and reveal cell-type–specific expression changes in type 2 diabetes. Genome research **27**(2), 208–222 (2017)

[7] Grün, D., Muraro, M.J., Boisset, J.-C., Wiebrands, K., Lyubimova, A., Dharmadhikari, G., Born, M., Van Es, J., Jansen, E., Clevers, H., *et al.*: De novo prediction of stem cell identity using single-cell transcriptome data. Cell stem cell **19**(2), 266–277 (2016)

[8] Vieira Braga, F.A., Kar, G., Berg, M., Carpaij, O.A., Polanski, K., Simon, L.M., Brouwer, S., Gomes, T., Hesse, L., Jiang, J., *et al.*: A cellular census of human lungs identifies novel cell states in health and in asthma. Nature medicine **25**(7), 1153–1163 (2019)

[9] Lotfollahi, M., Wolf, F.A., Theis, F.J.: scgen predicts single-cell perturbation responses. Nature methods **16**(8), 715–721 (2019)

[10] Xu, C., Lopez, R., Mehlman, E., Regier, J., Jordan, M.I., Yosef, N.: Probabilistic harmonization and annotation of single-cell transcriptomics data with deep generative models. Molecular systems biology **17**(1), 9620 (2021)

[11] Andreatta, M., Hérault, L., Gueguen, P., Gfeller, D., Berenstein, A.J., Carmona, S.J.: Semi-supervised integration of single-cell transcriptomics data. Nature Communications **15**(1), 872 (2024)

[12] Korsunsky, I., Millard, N., Fan, J., Slowikowski, K., Zhang, F., Wei, K., Baglaenko, Y., Brenner, M., Loh, P.-r., Raychaudhuri, S.: Fast, sensitive and accurate integration of single-cell data with harmony. Nature methods **16**(12), 1289–1296 (2019)

[13] Stuart, T., Butler, A., Hoffman, P., Hafemeister, C., Papalexi, E., Mauck, W.M., Hao, Y., Stoeckius, M., Smibert, P., Satija, R.: Comprehensive integration of single-cell data. Cell **177**(7), 1888–1902 (2019)

[14] Lopez, R., Regier, J., Cole, M.B., Jordan, M.I., Yosef, N.: Deep generative modeling for single-cell transcriptomics. Nature methods **15**(12), 1053–1058 (2018)

[15] He, C., Filippidis, P., Kleinstein, S.H., Guan, L.: Partially characterized topology guides reliable anchor-free scrna-integration. Communications Biology **8**(1), 561 (2025)
